# *Listeria monocytogenes* gene essentiality under laboratory conditions and during macrophage infection

**DOI:** 10.1101/2022.03.04.482958

**Authors:** Martin Fischer, Tim Engelgeh, Patricia Rothe, Stephan Fuchs, Andrea Thürmer, Sven Halbedel

## Abstract

The Gram-positive bacterium *Listeria monocytogenes* occurs widespread in the environment and infects humans when ingested along with contaminated food. Such infections are particularly dangerous for risk group patients, for whom they represent a life-threatening disease. To invent novel strategies to control contamination and disease, it is important to identify those cellular processes that maintain pathogen growth in- and outside the host. We here have applied transposon insertion sequencing (Tn-Seq) to *L. monocytogenes* for the identification of such processes on a genome-wide scale. Our approach classified 394 open reading frames as essential for growth under standard laboratory conditions and identified 42 further genes, which become additionally essential during intracellular growth in macrophages. Most essential genes encode components of the translation machinery, act in chromosome-related processes, cell division and biosynthesis of the cellular envelope. Several cofactor biosynthesis pathways and 29 genes with unknown functions were also essential, opening novel options for the development of anti-listerial drugs. Among the genes specifically required during intracellular growth were known virulence factors, genes compensating intracellular auxotrophies and several cell division genes. Our experiments also highlight the importance of PASTA kinase signalling, glycine metabolism and chromosome segregation for efficient intracellular growth of *L. monocytogenes*.

## INTRODUCTION

*Listeria monocytogenes* is foodborne pathogen causing life-threatening infections in humans and animals. The pathogen is transmitted from environmental sources via contaminated food and feed, can survive the passage through the gastrointestinal tract and may finally cross the gut epithelium to reach the bloodstream (1). Due to the ubiquitous presence of *L. monocytogenes* in the environment, food contaminations and thus periods of asymptomatic carriage in the gut are quite common. Approximately 5-10% of the population are considered to carry *L. monocytogenes* in the gastrointestinal tract and it is estimated that healthy adults experience two periods of asymptomatic *L. monocytogenes* carriage per year (2, 3). *L. monocytogenes* that crosses the epithelial barrier is rapidly cleared from the blood stream by macrophages and non-professional phagocytes in the liver and the spleen, which represent an important line of defence against the infection (4–6). *L. monocytogenes* may proliferate inside macrophages and hepatocytes and spread inside the liver parenchyma, even though this phase of infection is usually limited by innate immune defence and T-cell mediated cytotoxic mechanisms (6–8). In immunocompromised persons however, where an adequate immune response cannot be established, *L. monocytogenes* proliferation in the liver continues, facilitating systemic dissemination and translocation to other organs, primarily the brain and the uterus during pregnancy. Bacteraemia, cerebral and fetal infections are characteristically associated with high case fatality rates (9–12), and therefore, listeriosis is considered a major public health concern despite its low incidence (13).

*L. monocytogenes* is equipped with specific virulence factors to invade non-phagocytic cells in a phagocytosis-like process and to escape the phagosome after internalisation, to multiply within in their cytosol and to spread from cell to cell (6, 14). Six of the most important virulence factor genes (*prfA*, *plcA*, *hly*, *mpl*, *actA*, *plcB*) cluster in the *Listeria* pathogenicity island LIPI-1 (15). While these genes coordinate virulence factor expression (PrfA), act as hemolysin (Hly) and phospholipases (PlcA, PlcB) for penetration of host cell membranes or as nucleator of host actin polymerization for intracellular motility and cell to cell spread (ActA) (14), further proteins are required for intracellular proliferation. These proteins contribute to the biosynthesis of selected nucleotides (16, 17), aromatic amino acids (18) and menaquinones (19, 20) as well to the uptake of certain amino acids (21, 22) to compensate for auxotrophies occurring in the cytosol, or ensure efficient separation of daughter cells after completion cell division as a prerequisite for efficient intra- and intercellular spread (23, 24).

The identification of genes required for intracellular proliferation of *L. monocytogenes* and for growth of the bacterium in general is an important task to facilitate development of novel antilisterial therapies or to design more efficient strategies to prevent growth of the bacterium in food matrices. Up to now, essential *L. monocytogenes* genes were identified either in a stepwise gene-by-gene approach or by laborious screening of plasmid insertion mutant libraries (16, 25), but both techniques have important limitations. Tn-Seq is a technique that allows identification of genes which do not accept transposon insertions without loss of viability. Such genes can be collectively identified by massive parallel sequencing of transposon insertion libraries, which do not need to be separated into individual clones (26). Tn-Seq has been used to identify essential genes in a wide variety of bacteria. We here present, at least to our knowledge, the first application of this technique to *L. monocytogenes* and have used it to identify genes essential for growth of this important pathogen under standard laboratory conditions and during macrophage infection.

## MATERIAL AND METHODS

### Bacterial strains and growth conditions

All strains and plasmids used in this study are listed in Table 1. Strains of *L. monocytogenes* were cultivated in BHI broth or on BHI agar plates at 37°C. Antibiotics and supplements were added when required at the following concentrations: erythromycin (5 µg/ml), X-Gal (100 µg/ml) and IPTG (as indicated). *Escherichia coli* TOP10 was used as host for all cloning procedures (27). Ceftriaxone minimal inhibitory concentrations were determined using E-test strips with a ceftriaxone concentration range of 0.016 - 256 µg/ml (bestbion^dx^, Germany).

**Table 1:**
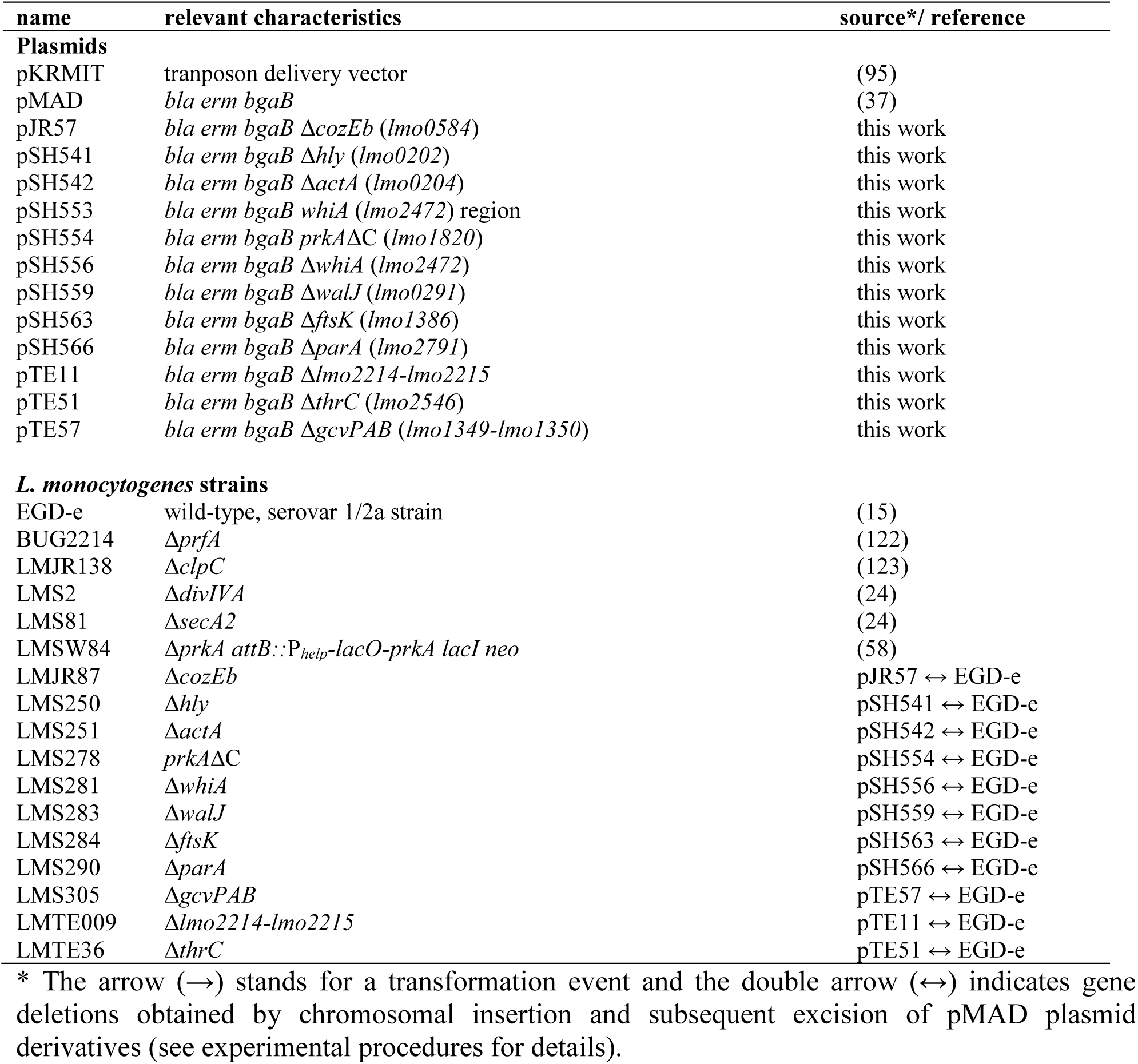
Plasmids and strains used in this study

### General methods, manipulation of DNA and oligonucleotide primers

Standard methods were used for transformation of *E. coli* and isolation of plasmid DNA (27). Transformation of *L. monocytogenes* was carried out as described by others (28). Restriction and ligation of DNA was performed according to the manufactureŕs instructions. All primer sequences are listed in Tab. 2.

**Table 2:**
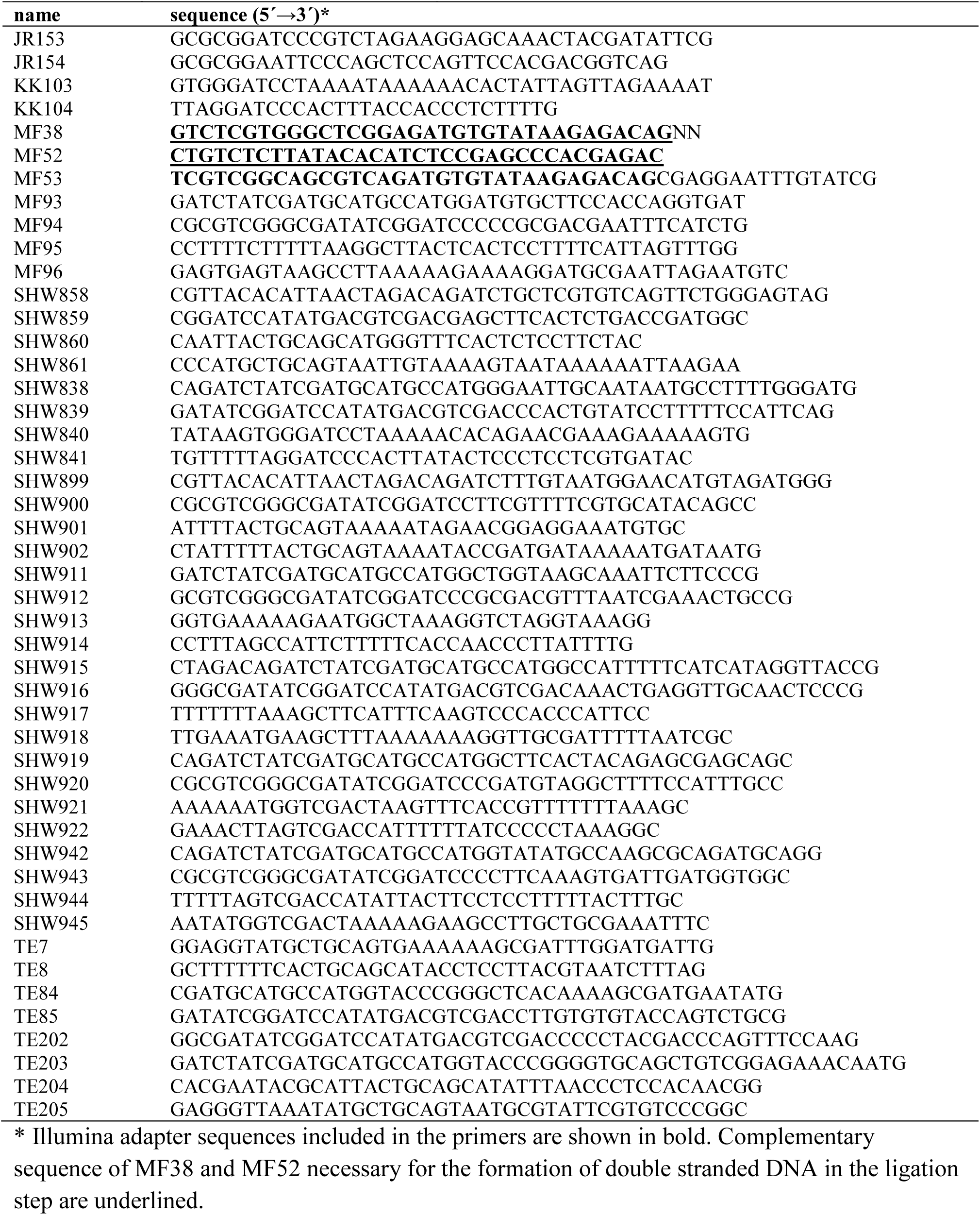
Oligonucleotides used in this study.

### Transposon library construction

The Tn-delivery vector pKRMIT plasmid was transformed in *L. monocytogenes* EGD-e. Transformants were selected on BHI agar plates containing 25 mg/l kanamycin and 5 mg/l spectinomycin after incubation at 30°C for 24 h. One clone was selected and grown overnight at 30°C in BHI broth containing both antibiotics. This culture was used to inoculate 25 ml BHI broth containing 25 mg/L kanamycin to a starting OD_600_ of 0.01 and then further incubated at 30°C until an OD_600_ of 0.1 was reached. Transposase expression and transposon integration was induced by increasing the cultivation temperature to 40°C and further incubation until an OD_600_ of 0.5. Aliquots were harvested, mixed with glycerol (50% final concentration) and stored at - 80°C. These aliquots were used to inoculate 25 ml of BHI broth containing kanamycin (25mg/l) to an OD of 0.1, which was grown at 37°C to stationary phase. This culture was diluted back and grown to stationary phase two more times to deplete clones that have lost the plasmid without transposition. At the end of the experiment, bacterial titers were determined on BHI agar plates and on BHI agar plates containing either kanamycin or spectinomycin to calculate the transposition frequency and the ratio of plasmid loss. 500 µl of the culture were centrifuged and genomic DNA was extracted for Tn insertion sequencing.

### Generation and sequencing of Tn-Seq amplicon libraries

Transposon library sample aliquots were used for DNA extraction by a phenol-chloroform isoamyl alcohol procedure (27). DNA pellets were resuspended in 200 µl 10 mM Tris/HCl pH 7.5 and DNA concentration was determined using Qubit dsDNA HS Assay.

3 µg of the isolated DNA were digested using 2 U MmeI at 37°C for 6 h followed by heat inactivation for 10 min at 80°C. Digestion was confirmed by agarose gel electrophoresis. 5 U of quick CIP calf intestinal phosphatase was directly added to the digestion mix and incubated for 1 h at 37°C. The reaction mix was then purified using a column-based PCR clean-up kit. Double stranded DNA adapters were obtained by mixing each 40 µl of the oligonucleotides MF38 and the 5’ and 3’ phosphorylated MF52 (both 100 µM) followed by denaturation at 99°C for 10 min. 3 µl of these adapters were added to the cleaned-up transposon library DNA and ligated using 5 U T4 DNA ligase. The produced fragments were purified using the Qiagen size selection kit and eluted in 20 µl 10 mM Tris/HCl pH7.5. Sequencing adapters were attached to the fragments via PCR. For this purpose, 8 µl of the eluate were used as template in an 80 µl PCR reaction with MF38 and MF53 as the primers. The size of the PCR products was controlled using agarose gel electrophoresis and the products were purified from the gel. The adapter-linked PCR products were used as a template to introduce multiplex identifiers for pooled sequencing. PCR was performed with following conditions: 2.5 µl Template (3 ng/µl), 2.5 µl Nextera XT index primer (each N7 and S5, Illumina) and 12.5 µl 2x KAPA HiFi HotStart ReadyMix (Roche) and 5 µl H2O using a PCR program with these settings: 95°C 3 min, 8 cycles (each 30 sec) of 95°C, 55°C, 72°C, 72°C 5 min and storage at 4°C until bead-clean up. Sequencing was performed in a 2 x 150 bp paired end run on a NextSeq550 instrument (Illumina) using v2.5 midout chemistry.

### Bioinformatic processing of sequence reads and gene essentiality prediction

Only raw reads containing the terminal fraction of the transposon were accepted for further analysis. The sequence region of the inverted repeat was afterwards removed using cutadapt (29) and 13 bases following the inverted repeat were kept for determination of the insertion locus. The insertion locus was determined by back-mapping the trimmed sequences against the *L. monocytogenes* EGD-e genome (30) using bowtie with the options “-S -v 0 -n 0 -p 20 -l 13 -m 1” allowing no mismatch within the alignment and only unique insertion sites to be counted (31). The resulting sam file was used for the analysis in Tn-Seq Explorer version 1.5 (32). For the determination of zbar values, the sam file was further transformed to wiggle file format using an in-house script and analysed using the software TRANSIT version 3.1.0 (33).

Tn-Seq Explorer and TRANSIT were used for the prediction of essentiality and determination of prediction accuracy (zbar). For Tn-Seq Explorer analysis, sequencing run data in the form of the sam file and the genbank file of *L.monocytogenes* EGDe (NC_003210.1) were uploaded to TnSeq-Explorer. Insertion loci were excluded if they were covered by less than five reads mapping onto a them. Repetitive hits of one insertion site were counted as one unique insertion, so that loci potentially preferred during PCR-amplification would not compromise further analysis. Essential genes were defined as genes with an insertion density lower 0.01. The insertion density is defined as the number of unique insertions divided by the number of bases of the gene (32). Identified essential genes were visually confirmed using the Artemis comparison tool (34).

### Premature stop codon analysis

Assembled genome sequences of all of *Listeria monocytogenes* isolates that were available at the NCBI pathogen detection pipeline server (https://www.ncbi.nlm.nih.gov/pathogens) in December 2019 (n=27118) were downloaded. All fasta files were imported in SepSphere (Ridom) to perform core genome and accessory genome multi locus sequence typing (cgMLST/agMLST) by automated allele submission to cgMLST.org (35). Afterwards, all allele sequences for all targets of the *L. monocytogenes* cgMLST (n=1701) and agMLST schemes (n=1158) deposited at cgMLST.org were downloaded and alleles with premature stop codons between 5-80% relative to the start of the open reading frame were determined for each target using an in-house script. Frequency of occurrence of such alleles among the 27118 genomes was quantified.

### Construction of plasmids and strains

Plasmid pSH541 was generated for deletion of *hly*. For its construction, *hly* up- and downstream fragments were amplified using the primers SHW858/SHW860 and SHW861/859, respectively, fused together by overlapping extension (SOE)-PCR and inserted into pMAD by restriction free (RF) cloning (36).

Plasmid pSH542 was generated for deletion of *actA*. Fragments up- and downstream of *actA* were amplified by PCR using the oligonucleotides SHW838/SHW841 and SHW840/SHW839, respectively. Both fragments were spliced together by SOE-PCR and then inserted into pMAD using RF cloning.

Plasmid pSH554, facilitating deletion of the *prkA* C-terminus, was obtained by amplification of fragments up- and downstream of the *prkA* region to be deleted using the primer pairs SHW899/SHW902 and SHW901/SHW900, respectively, which were then fused together by SOE-PCR and inserted into pMAD by RF cloning.

A chromosomal fragment containing *whiA* together with its up- and downstream regions was amplified using the primers MF93/MF94 and inserted into pMAD by RF cloning, which resulted in plasmid pSH553. The *whiA* gene of this plasmid was then removed in a PCR with MF95/MF96 as the primers, yielding the *whiA* deletion plasmid pSH556.

Plasmid pJR57 was constructed for deletion of *cozEb* (*lmo0584*). To this end, chromosomal regions up- and downstream of *cozEb* were amplified using JR153/KK104 and KK103/JR154, respectively, as the primers and spliced together by SOE-PCR. The resulting fragment was then cloned into pMAD using EcoRI/BamHI.

Likewise, plasmid pSH559 was constructed for deletion of *walJ* (*lmo0291*). Regions flanking the *walJ* gene were amplified using the oligonucleotides SHW919/SHW922 and SHW921/SHW920 and spliced together by SOE-PCR. Finally, the resulting fragment was inserted into pMAD by RF cloning For construction of plasmid pSH563, designed for deletion of *ftsK* (*lmo1386*), fragments up- and downstream to *ftsK* were amplified using SHW915/SHW917 and SHW916/SHW918, respectively, as the primers. Both fragments were spliced together by SOE-PCR and inserted into pMAD by RF cloning.

Fragments up- and downstream to *parA* (*lmo2791*) were amplified by PCR using oligonucleotides SHW911/SHW914 and SHW913/SHW912, respectively, and then fused together by SOE-PCR. The resulting fragment was inserted into pMAD by RF cloning, yielding plasmid pSH566.

For construction of plasmid pTE11, fragments up- and downstream of *lmo2214-lmo2215* were amplified by PCR using TE8/TE85 and TE84/TE7, respectively, fused together by SOE-PCR and inserted into pMAD by RF cloning.

Plasmid pTE51 was designed for deletion of *thrC* (*lmo2546*). Fragments up- and downstream of *thrC* were amplified by PCR using TE203/TE204 and TE205/TE202, respectively, as the primers. The two fragments were spliced together in a SOE-PCR and then inserted into pMAD by RF cloning.

Plasmid pTE57, constructed for deletion of *gcvPAB* (*lmo1349-lmo1350*), was generated similarly. First, regions up- and downstream of *gcvPAB* were amplified by PCR using the oligonucleotides SHW942/SHW944 and SHW945/SHW943, respectively. In a second step, both fragments were joined by SOE-PCR and the resulting fragment was inserted into pMAD by RF cloning.

Derivatives of pMAD plasmids were transformed into the *L. monocytogenes* recipients and genes were deleted as described elsewhere (37). All gene deletions were confirmed by PCR.

### Isolation of cellular proteins and Western blotting

Cells were grown in BHI broth at 37°C to an optical density of OD_600_=1.0 and harvested by centrifugation. Cell pellets were washed with ZAP buffer (10 mM Tris.HCl pH7.5, 200 mM NaCl), resuspended in 1 ml ZAP buffer to which 1 mM PMSF was added and disrupted by sonication. Cellular debris was removed by centrifugation and the supernatant was considered to contain total cellular proteins. Sample aliquots were separated by standard SDS polyacrylamide gel electrophoresis. Gels were transferred onto positively charged polyvinylidene fluoride membranes by semi-dry transfer. DivIVA and MurA were immune-stained using polyclonal rabbit antisera originally generated against *B. subtilis* DivIVA (38) and MurAA (39) as the primary antibodies and an anti-rabbit immunoglobulin G conjugated to horseradish peroxidase as the secondary one. The ECL chemiluminescence detection system (Thermo Scientific) was used for detection of the peroxidase conjugates on the PVDF membrane in a chemiluminescence imager.

### Infection experiments

Infection experiments with the transposon library were carried out in J774.A1 mouse ascites macrophages (ATCC) similar to a previously published protocol (40) but with some modifications. 10^7^ macrophages were seeded in 500 ml coated cultivation flasks (VWR) containing 75 ml DMEM + 10% FCS and incubated for 48 h at 37°C and 5% CO2. For bacterial infection, an overnight culture of the transposon library incubated at 37°C at 250 rpm in BHI was adjusted to an OD_600_ of 0.001 in DMEM. J774.A1 macrophages were infected with 10^6^ bacterial cells, resulting in a MOI=0.1. After 1 h of infection incubation, the media was removed from the cell culture, cells were briefly washed with 10 ml room temperature DMEM + 10% FCS and further incubated for 1 h in 50 ml DMEM + 10% FCS + 40 mg/l gentamicin to eliminate free bacterial cells. After 1 h, the media was removed and 75 ml of DMEM + 10% FCS + 10 mg/l gentamicin was added to the cell culture which was then further incubated for 24 h. At the end of the incubation, cells were harvested in 10 ml cold Triton X-100 in PBS, 50 µl of which was used for dilution series on BHI-agar plates to determine the number of viable bacterial cells recovered from the infected macrophages, while the rest of the cells were centrifuged at 5000 x g for 15 min at 4°C, decanted and stored for DNA extraction at -20°C.

J774.A1 macrophage infection experiments and plaque formation assays with 3T3 mouse fibroblasts for analysis of individual deletion mutants were essentially carried out as described earlier (40).

## RESULTS

### Tn-Seq based identification of genes required for growth under laboratory conditions

A highly saturated *mariner* transposon library was generated in the *L. monocytogenes* reference strain EGD-e with ∼79.000 individual clones. Thus, more than 30% of the 216,636 TA sites (0.07 TA sites per bp), which serve as possible transposon insertion sites were factually hit by a transposon. The library was cultivated in BHI broth at 37°C for 36 h (∼45 generations) and chromosomal DNA was isolated. Massive parallel sequencing of the transposon DNA junctions (Tn-Seq) (26) was used for identification of all Tn insertion sites. This uncovered 66,708 unique insertion sites confirming that 30.8% of all possible insertion sites were hit. Insertion sites were uniformly scattered around the chromosome as a consequence of the uniform distribution of TA sites (Fig. 1A). For the identification of essential genes, transposon insertions in the N-terminal 5% of the open reading frames (ORF) and C-terminal 20% of the ORFs were neglected and insertions were only counted if they were replicated at least 3 times. In addition, genes with an insertion density as defined by Langridge (41) below 0.01 (average insertion density in the dataset 0.32) were added, summing up to 394 essential genes (Tab. S1). An assignment of functional categories showed that most of these genes encode components of the translation machinery, act in replication, segregation and maintenance of the chromosome or are involved in biosynthesis of the cell envelope (Fig. 1B). The function of 29 essential genes is not known (Tab. S1). Using previously published RNA-Seq data (42), we found that the essential genes are on average fivefold stronger expressed and not a single unexpressed gene was among them (Fig. 1C), which is in good agreement with their requirement for growth. Furthermore, the set of 394 genes identified here largely overlaps with the 258 *Bacillus subtilis* essential genes currently listed on Subtiwiki (43) (Fig. 1C). The 46 essential genes specific to *B. subtilis* either do not have a homolog in *L. monocytogenes* (n=13), are non-essential in *L. monocytogenes* (n=22) or are considered non-essential by the Tn-Seq-Explorer algorithm because Tn insertions are present in certain domains only (n=3) or just above the cut-off (n=8). We categorize these latter 8 genes as potentially essential (Table S1).

**Figure 1:**
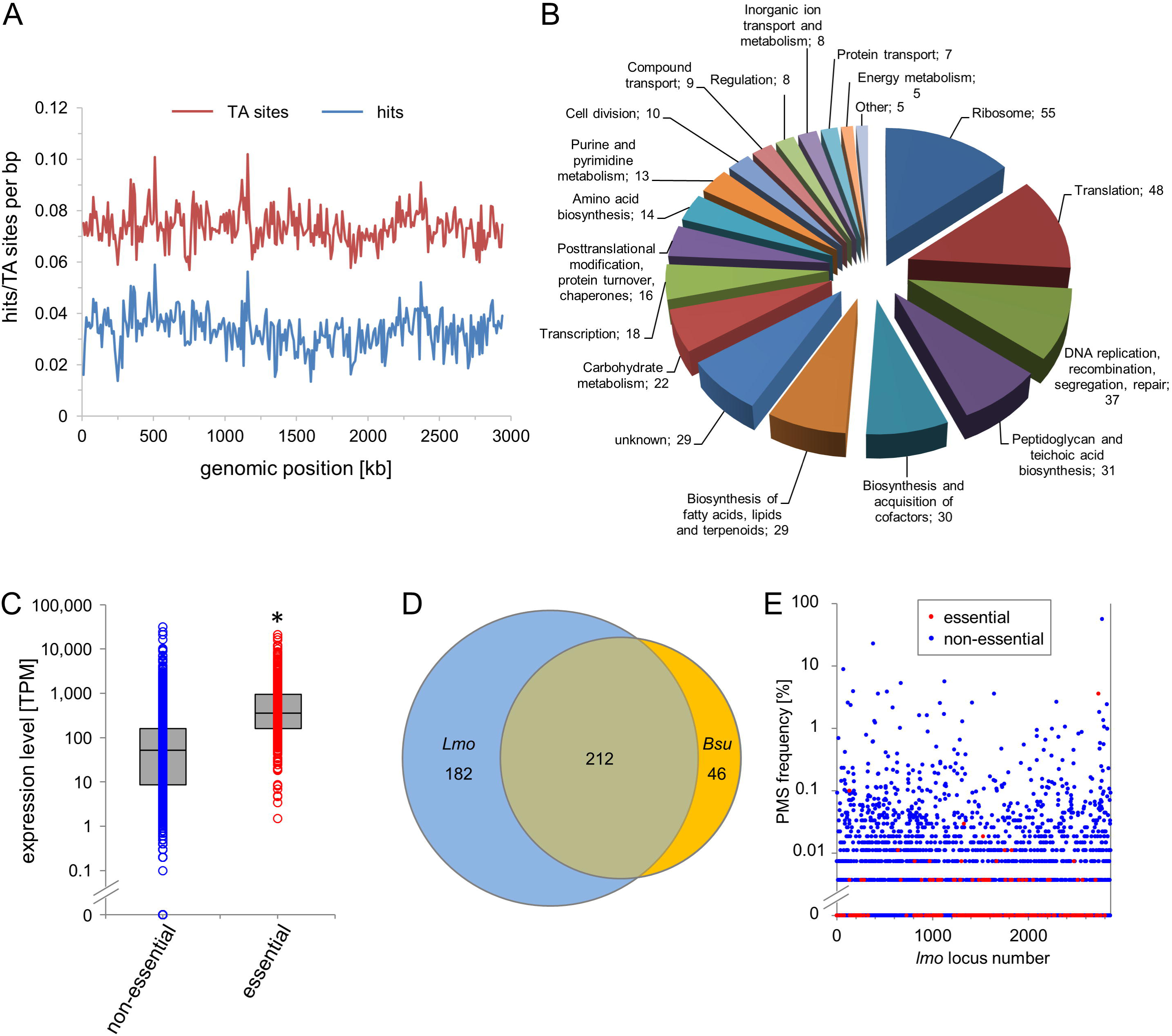
Identification of essential *L. monocytogenes* genes by Tn-Seq. (A) Distribution of TA sites and Tn insertion along the *L. monocytogenes* EGD-e genome. TA sites/hits per bp were calculated using a coverage window of 10,000 bp. (B) Functional classes of the 394 essential genes of *L. monocytogenes* EGD-e. (C) Expression levels of the 394 essential genes in comparison to non-essential genes according to a previously published RNA-Seq experiment with *L. monocytogenes* EGD-e under standard growth conditions (42). The asterisk indicates statistical significance (*t*-test, *P*<0.0001). (D) Venn diagram illustrating the intersection between the essential genes of the *L. monocytogenes* EGD-e (n=394) and *B. subtilis* 168 (n=258). (E) Frequency of naturally occurring premature stop codons (PMS) in the each of *L. monocytogenes* EGD-e genes as observed in a dataset of 27,118 *L. monocytogenes* genomes from clinical and environmental isolates that were downloaded from the NCBI server.

182 genes were found to be essential in *L. monocytogenes* without being annotated as essential in *B. subtilis*. 22 of these genes had no homolog in *B. subtilis*. For the remaining 160 genes several explanations seem possible: (i) Those genes become specifically essential at increased temperature used during transposition (such as *gpsB, clpP, dnaK* or *lmo1450*) (44–47), (ii) a severe growth defect results from their deletion without being essential *per sé* (such as *ackA, ltaS, oppF, ptsI, pycA, sipZ, sodA* among others) (48–54), or (iii) they are indeed essential in *L. monocytogenes* but not in *B. subtilis* (*e. g. ezrA*, *cdaA*, *ftsA*, *pbpB1*, *prpC*) (55–58). This latter class of genes can be explained by a reduced number of potential back-up enzymes that can be encoded in an organism with a smaller genome. A minority of 8 genes was categorized as essential by the Tn-explorer algorithm even though previous work reported their deletion without adverse effects on growth (*e. g. codY, cshA, cspD, fvrA, recA, relA, secDF*) (59–63) (Fig. S1). The reason for these potentially false-positive identifications is currently not known (see discussion).

### Genes with naturally occurring premature stop codons

Genome sequences of *L. monocytogenes* isolates from environmental, food and clinical samples currently are becoming available in high number in the course of pathogen surveillance programs. Allelic information of many of these genomes is stored on genomic subtyping servers such as cgMLST.org (35). We extracted the allelic variations known in the *L. monocytogenes* population for all 2857 EGD-e genes from the cgMLST.org server and identified alleles containing premature stop codons (PMS) located within 5% and 80% of their sequence relative to their start. Next, we counted the number of genomes that carried such PMSs for each of the 2857 EGD-e open reading frames using a collection of 27,118 *L. monocytogenes* genomes from clinical and environmental isolates available at the NCBI pathogen detection pipeline at the time of analysis (64). This showed that no naturally occurring premature stop codons are known for 1424 ORFs, an observation that can be interpreted as functional gene essentiality under real life conditions. 348 (87%) of the 394 essential and 8 potentially essential genes did not contain PMSs and 42 (10%) of them had a PMS only once (Fig. 1E). Thus, the majority of all laboratory confirmed essential genes also does not lose its functionality outside the lab. In contrast, higher PMS frequencies were observed among the non-essential genes (Tab. S2). Several genes, particularly those coding for extracellular proteins (among them: *inlA*) showed PMS frequencies above 1%, which likely reflects natural antigenic variation. Taken together, our experimental assignment of gene essentiality through Tn-Seq inside the lab is in congruence with the observed frequency of gene functionality loss through spontaneous allelic variations leading to PMSs under real life conditions.

### Gene essentiality in central carbon metabolism

*L. monocytogenes* generates ATP from glycolytic metabolism of hexoses or glycerol. In good agreement with the redundancy of phosphotransferase systems (PTS) or glycerol uptake facilitators, none of the uptake systems for any of these carbon sources was essential. In contrast, the genes for the complete glycolytic reaction chain were essential (Fig. 2, Fig. 3) with the exception of *pykA*, coding for pyruvate kinase (Fig. 3). Presumably, pyruvate can be synthesized from phosphoenolpyruvate by pyruvate phosphate dikinase (65) or by PTS enzyme I proteins when *pykA* is inactivated as observed in *B. subtilis* (66). Pyruvate is further catabolized to lactate, acetate, acetyl-CoA, oxaloacetate or acetoin (52, 67, 68). The *ackA1* gene encoding acetate kinase for acetate formation and several pyruvate dehydrogenase component genes for acetyl- CoA formation are essential. With pyruvate oxidase (*lmo0722*) and phosphotransacetylase (*lmo2103*), two alternatives for formation of acetyl phosphate exist, explaining why both genes are dispensable despite essentiality of *ackA1* (Fig. 3). Likewise, *ldh*, encoding lactate dehydrogenase and its three paralogs were dispensable, as were the *alsS* and *alsD* genes required for acetoin formation. Pyruvate carboxylase PycA generates oxaloacetate for biosynthesis of several amino acids and, consequently, the *pycA* gene was essential.

**Figure 2:**
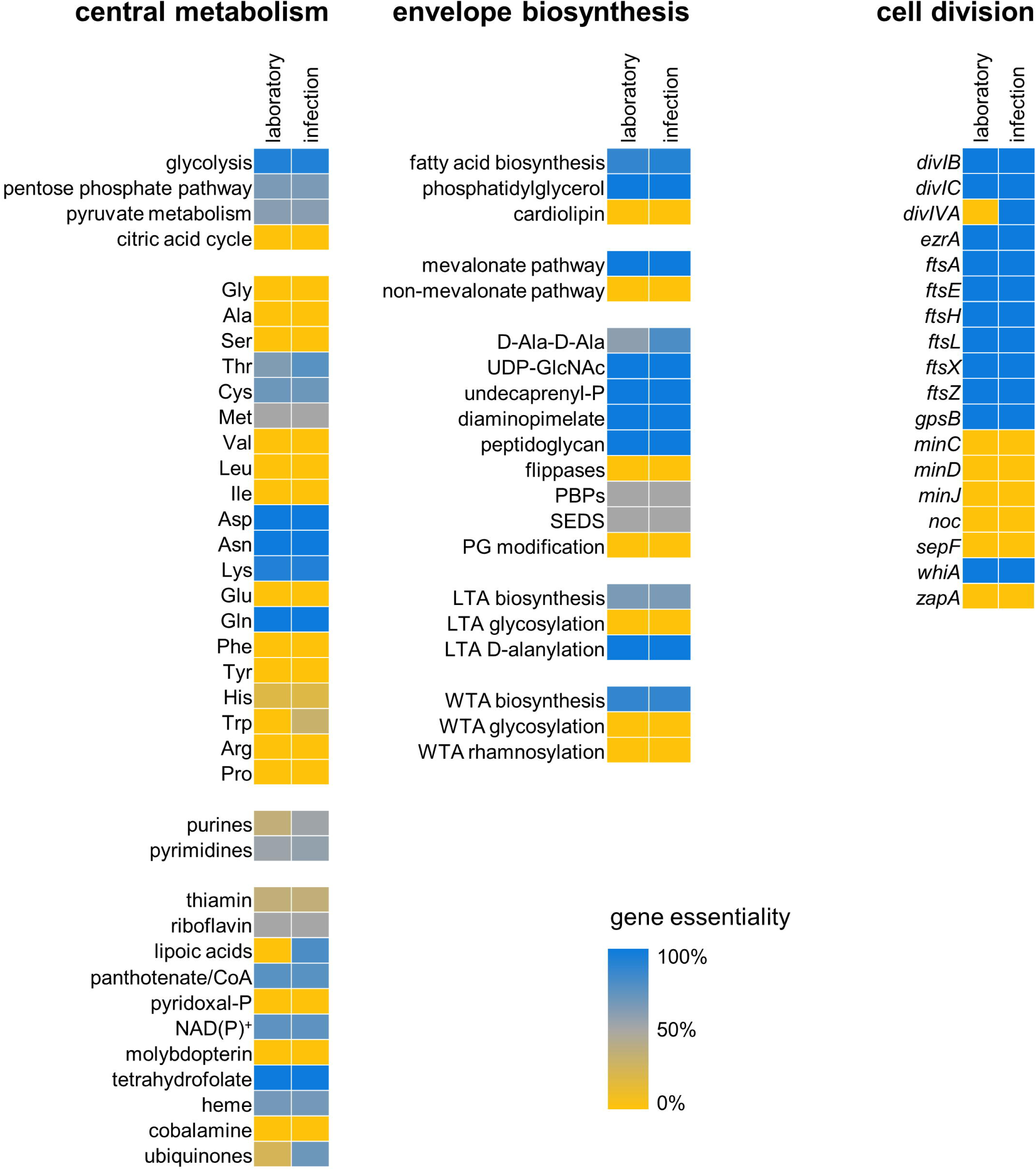
Gene essentiality in central metabolism, envelope biosynthesis and cell division. Heat maps illustrating the essentiality of selected genes and pathways. Where genes were aggregated according to their pathways, gene essentiality is expressed as the number of essential divided by the number of all genes in this particular pathway. Gene essentiality is shown for standard growth conditions (laboratory) and during growth in J774 macrophages (infection).

**Figure 3:**
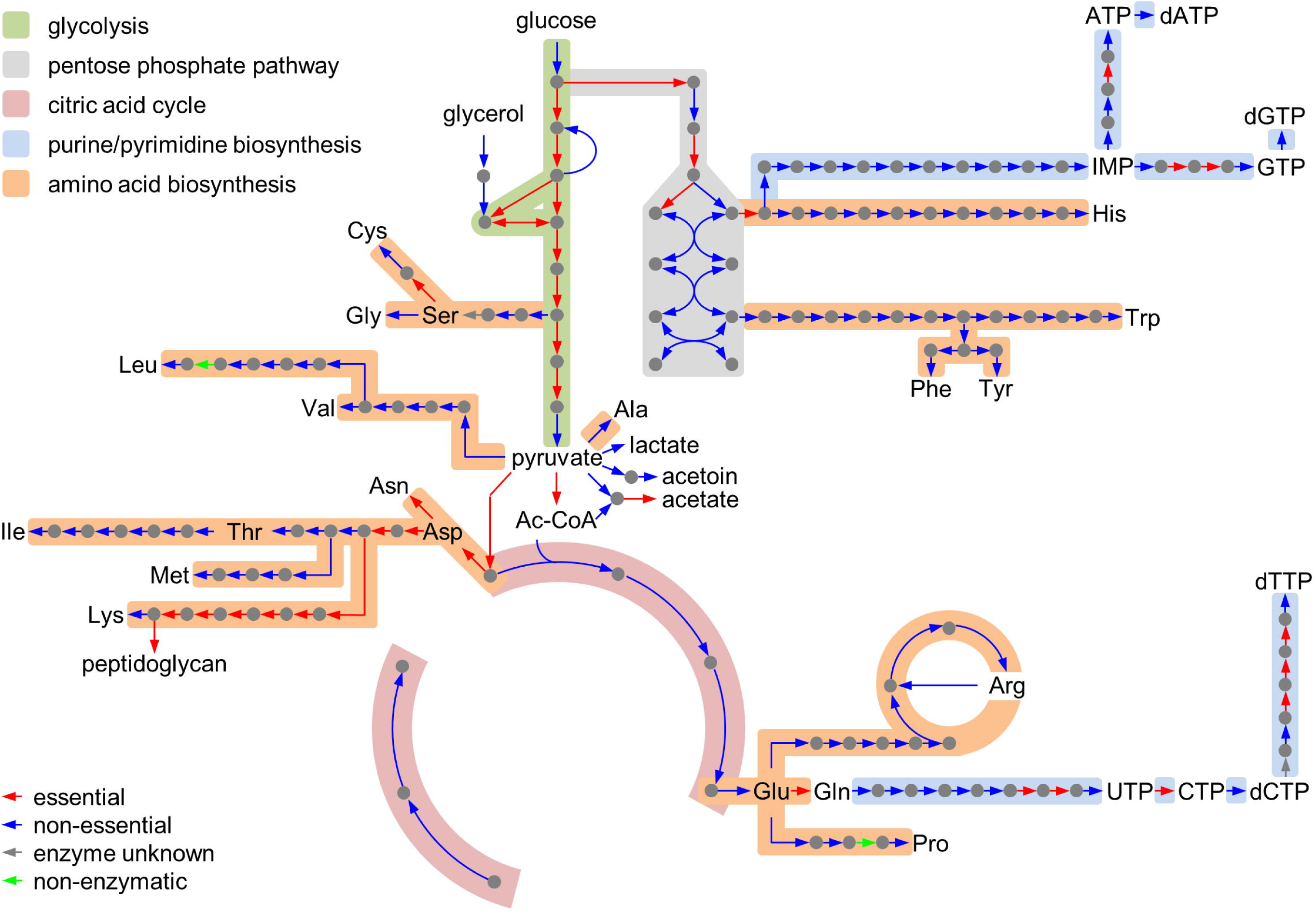
Gene essentiality in central carbon and nitrogen metabolism. Scheme illustrating the metabolic pathways in central carbon metabolism, the biosynthesis of nucleobases and their corresponding nucleotides and biosynthesis of amino acids according to the KEGG database (https://www.genome.jp/kegg-bin/show_organism?org=T00066). Reactions are colored according to the essentiality of the corresponding genes.

The tricarboxylic acid (TCA) cycle is incomplete in *L. monocytogenes* (69), and in line with this, all TCA cycle genes were dispensable (Fig. 2, Fig. 3). In contrast, selected genes for the upper oxidative part of the pentose phosphate pathway (PPP) were not found to be disrupted, whereas the genes in the lower non-oxidative part could largely be inactivated (Fig. 3). PPP dispensability is in good agreement with results showing that pentitols can be taken up for pentose formation (70). Moreover, the enzymatic reactions in the lower non-oxidative part are encoded by sets of highly redundant genes. The only essential enzyme in this part of the pathway is *lmo1818*, which encodes one out of six paralogous ribulose-phosphate 3-epimerase genes. Interestingly, *lmo1818* was the only one expressed under standard growth conditions in a previous transcriptome analysis (42).

All steps in the biosynthesis of fatty acids from acetyl-CoA are mediated by essential enzymes (Fig. 2, Fig. S2). One exception is enoyl-acyl carrier protein reductase, which occurs in four isoforms (FabI, FabL, FabK1 and FabK2) (71). The *fabI* and *fabL* genes are essential, whereas *fabK1* and *fabK2* are individually dispensable. Presumably their simultaneous deletion is not tolerated. Taken together, glycolytic glucose conversion to acetyl-CoA for fatty acid biosynthesis or to acetate for energy conservation represent the essential backbone of *L. monocytogenes* carbon catabolism (Fig. 2, Fig. 3).

### Biosynthesis and acquisition of amino acids, purines and pyrimidines

The biosynthetic pathways for all 20 proteinogenic amino acids are complete, but only some genes were essential. Among them were the genes for aspartate amino transferase AspB (*lmo1897*), asparagine synthase AnsB (*lmo1663*), glutamine synthetase GlnA (*lmo1299*) and serine acetyltransferase CysE (*lmo0238*, Fig. 2, Fig. 3), which fits with the observation that *L. monocytogenes* could grow in defined minimal medium not containing glutamine, asparagine or aspartate (72). All genes required for lysine biosynthesis except the last gene in this pathway (*lysA*) were also essential (Fig. 3). LysA catalyzes lysine production from diaminopimelate. This essentiality pattern shows that lysine can be acquired from external resources, whereas diaminopimelate is essential for peptidoglycan (PG) biosynthesis and must be synthesized. Two peptide transport systems or their components turned out to be essential: The *ctaP* (*lmo0135*)- *lmo0136*-*lmo0137* cysteine transporter and the *oppD* and *oppF* ATPase subunits of the *oppABCDF* oligopeptide transporter (21). This suggests that uptake is the preferred mode of amino acid acquisition in rich medium, as previously suggested by metabolomics (73). *L. monocytogenes* encodes all genes necessary for *de novo* biosynthesis of purines and pyrimidines (15) (Fig. 3) and additionally contains several genes for nucleobase uptake (*lmo0456*, *lmo0573*, *lmo1839/pyrP*, *lmo1845*, *lmo1884*, *lmo2254*). Since biosynthesis and uptake can mutually replace each other, the genes for these transporters and all genes required for nucleobase biosynthesis were dispensable. In contrast, many of the genes required for generation and interconversion of nucleotides are essential (Fig. 2). Thus, our gene essentiality data reveal a certain flexibility in the listerial N-metabolism which likely can switch between synthesis and uptake depending on the nutrient supply.

### Cofactors and vitamins

*L. monocytogenes* requires riboflavin, thiamine, biotin and lipoic acid to grow in minimal medium (72, 74). Consequently, their biosynthetic pathways are incomplete (15) and the few remaining genes are non-essential, whereas the riboflavin transporter (*lmo1945*) could not be disrupted (75). Interestingly, the only known thiamine (*lmo1429*) (76) and biotin uptake genes (*lmo0598*) (77) were not essential, suggesting the presence of alternative uptake routes. The *pdxST* genes encoding pyridoxal phosphate synthase were also not essential (Fig. 2), likely because pyridoxal can be imported via an unknown transport mechanism (78, 79).

The entire gene set for the *de novo* formation of NAD(P)^+^ from aspartate and nicotinate is present, but only *nadD*, *nadE*, and *nadF* (*lmo0968*) for the terminal reactions are essential (Fig. S3). Presumably, nicotinate can be taken up by an unknown transporter, which also would explain the essentiality of *pcnB* (*lmo1092*) that funnels nicotinate into this pathway just ahead the NadDEF-mediated steps.

Two pathways feed pantothenate biosynthesis, one starting from pyruvate (*ilvB*, *ilvH*, *ilvC*, *ilvD*) and the second from valine (*lmo0978*). Genes for both pathways are non-essential up to the point where they merge and the genes downstream are essential (*panB*, *lmo2046*, *panC, panD*) as were the genes necessary for coenzyme A formation from pantothenate (*lmo1825*, *coaD*, *coaE, acpS, acpP*). The *lmo0221* and *lmo0922* encoding two alternative pantothenate kinases were a notable exception (Fig. S3).

The eight genes for molybdopterin biosynthesis from GTP are complete in EGD-e (*lmo1038*, *lmo01042*, *lmo1043*, *lmo1044*, *lmo1045*, *moaC*, *moaA*, *lmo1048*), but none of them is essential, suggesting alternative routes for molybdopterin acquisition (Fig. S3). In contrast, all genes for biosynthesis of tetrahydrofolate (THF) from GTP (*folE*, *folA*, *folK*, *sul*, *folC*, *lmo1873*) and THF regeneration (*fmt*, *folD*, *thyA*) are essential (Fig. S3), reflecting the susceptibility of *L. monocytogenes* towards folate biosynthesis inhibitors such as cotrimoxazol.

The genes for heme, vitamin B12 and menaquinone biosynthesis were completely present. A significant number of heme biosynthesis genes was essential (Fig. S4), despite the presence of heme transporters (80). In contrast, genes for vitamin B12 and menaquinone biosynthesis were largely dispensable (Fig. 2, Fig. S4).

Taken together, essential enzymes acting in biosynthesis of pantothenate/CoA, tetrahydrofolate or NAD(P)^+^ could be the promising targets for identification of novel compounds inhibiting growth of *L. monocytogenes*.

### Envelope biosynthesis and cell division

More stringent essentiality patterns are observed in the pathways for biosynthesis of the cellular envelope (Fig. 2). Genes for the cytosolic steps of PG biosynthesis up to lipid II (*glmSMU*, *murABCDEF*, *mraY*, *murG*) are collectively essential, as well as the genes for diaminopimelate production from aspartate and generation of D-Ala-D-Ala from D-alanine (Fig. 2, Fig. S2). The two enzymes for generation of undecaprenyl-P from farnesyl-PP are also essential (*lmo1315*, *lmo2017*), but from the two pathways for isoprenoid biosynthesis, only the mevalonate pathway genes were not hit by transposons, whereas all MEP/DOXP pathway genes could be inactivated (Fig. 2, Fig. S2). *Listeria* encodes two potential MurJ-like flippases for lipid II transport (*lmo1624*, *lmo1625*) and both could be disrupted. Presumably, both enzymes can mutually replace each other. Moreover, only one (*ftsW1*) out of six SEDS-type transglycosylases and two (*pbpB1*, *pbpB2*) out of five high molecular weight penicillin binding proteins were essential (Fig. S2), as shown in previous gene deletion experiments (57, 81). This is explained by a high degree of redundancy among the enzymes ensuring the extracellular steps of PG biosynthesis. Similarly, the genes participating in phospholipid biosynthesis were generally essential, but cardiolipin synthase was not required (Fig. S2). Also, the genes contributing to formation of wall teichoic acids (WTAs) were not inactivated by Tn insertions, with the exception of the two *tagO* genes (*lmo0959* and *lmo2519*), two paralogs for the first enzymatic step, which also can mutually compensate each other (82). Interestingly, the *dltABCD* genes for WTA decoration with D-alanine were all essential, whereas the *rmlTACBD* and *lmo2550, gtcA* and *lmo1079* genes for rhamnosylation (83) and glycosylation of WTAs (84), respectively, as well as several key enzymes in lipoteichoic acid biosynthesis were dispensable (Fig. S2). This indicates a functional hierarchy in the importance of the different types of teichoic acids and their modification systems, but also shows that generation of membranes, PG and WTAs altogether are generally critical for listerial viability. Likewise, most core divisome components were essential, however, genes with auxiliary function in divisome assembly and turnover (*divIVA*, *minCD*, *minJ*, *sepF* and *zapA*) were found inactivated under standard laboratory conditions (Fig. 2). The *lmo2472* gene, encoding a homologue of the *B. subtilis* cell division gene *whiA* was an interesting exception, since a *whiA* deletion only gave a mild phenotype in *B. subtilis* (85), whereas *lmo2472* was essential according to Tn-Seq. Despite this, a mutant lacking this gene could be generated and was viable at 37°C, but unable to grow at 42°C (Fig. S5). This illustrates a limitation associated with the chosen Tn delivery system, which relies on an incubation step at 40°C and thus also counter-selects heat-sensitive mutants.

### Implications for PASTA kinase signalling

We previously showed that *prkA*, encoding a eukaryotic-like serine/threonine protein kinase important for regulation of PG biosynthesis, is essential in *L. monocytogenes* EGD-e (58, 86). Among the substrates of this kinase are YvcK (*lmo2473*) and ReoM (*lmo1503*), both contributing to PG biosynthesis (58, 86–88). The precise role of YvcK is not clear, but the *yvcK* gene is encoded in the same operon next to *whiA* (30), herein identified as an essential gene at increased temperature. ReoM controls ClpCP-dependent degradation of MurA, which catalyses the first committed step in PG biosynthesis (58). In contrast to our previous results, *prkA* was hit by Tn-insertions. However, all Tn insertions in *prkA* clustered in the C-terminal half of the gene and thus separated the extracellular PASTA domains from the cytoplasmic kinase domain (Fig. 4A). To confirm that the *prkA* kinase domain and not the PASTA domains are required for viability, we tried to remove the *prkA* C-terminus from the chromosome. A viable mutant lacking the PASTA domains (*prkA*ΔC) could be generated and grew with wild type kinetics under standard laboratory conditions (Fig. 4B). The *prkA*ΔC strain was susceptible to ceftriaxone (Fig. 4C), suggesting perturbation of PG biosynthesis. MurA levels were not reduced as previously shown for a PrkA-depleted cells (58). Instead, the MurA level in the *prkA*ΔC mutant accumulated slightly above the wild type level. (Fig. 4D). Since situations, in which ReoM cannot be phosphorylated, lead to rapid MurA degradation and thus are lethal (58), this suggests that the isolated PrkA kinase domain is active in ReoM phosphorylation *in vivo*, which is in good agreement with previous *in vitro* data (58). Apparently, the kinase domain provides the essential enzyme function, whereas the PASTA domains are dispensable, suggesting that the PASTA domains exert a regulatory effect on the PrkA kinase. Remarkably, this effect becomes important during infection, as the *prkA*ΔC strain shows a significant delay in intracellular replication (Fig. 4E) that cannot be explained by a general growth defect (Fig. 4B).

**Figure 4:**
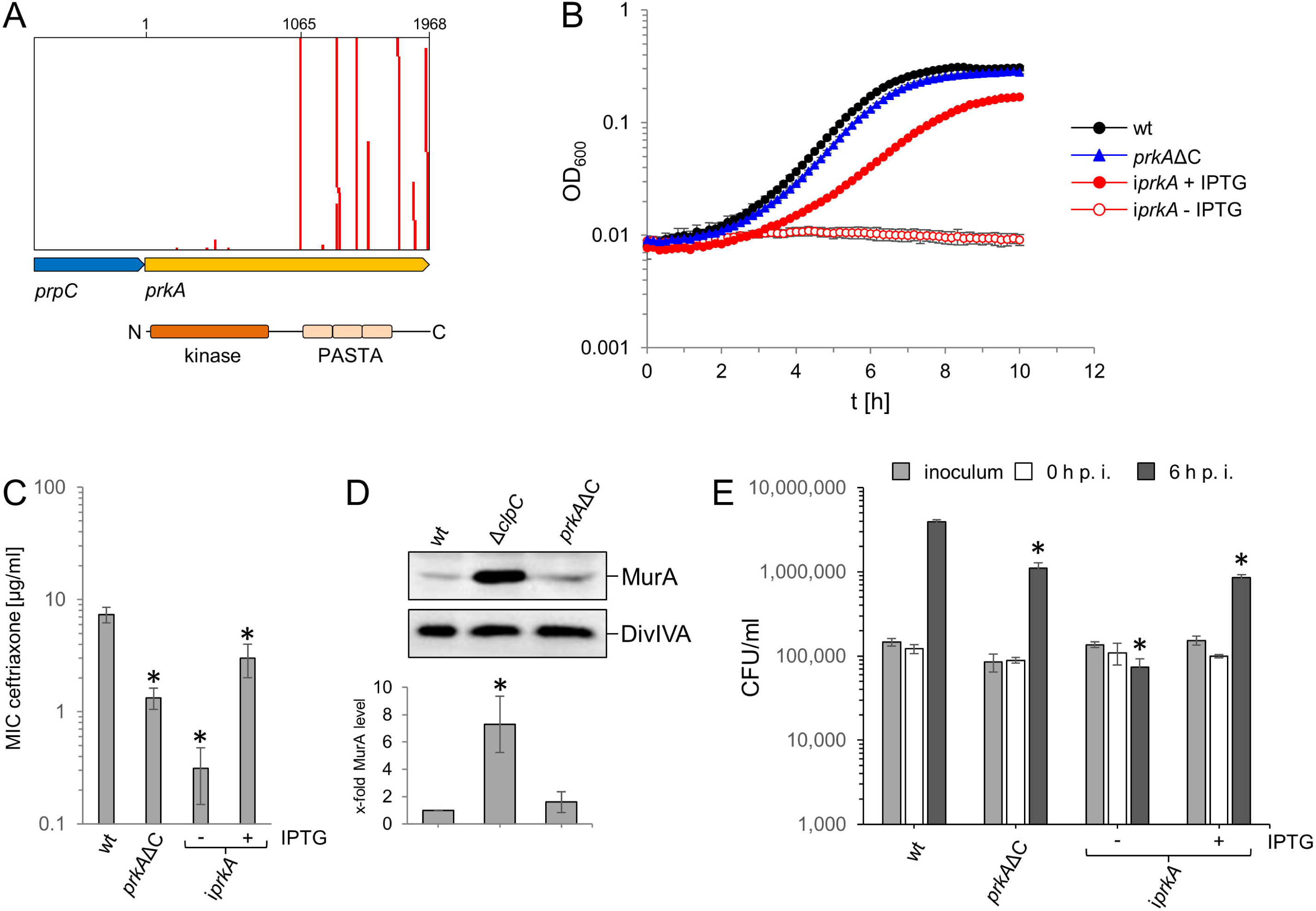
Dispensability of the *prkA* PASTA domains for *L. monocytogenes* viability. (A) Tn insertions at the *prkA* locus. Numbers refer to nucleotides in the *prkA* gene. The arrangement of PrkA domains is given below. (B) Viability of a *L. monocytogenes prkA*ΔC mutant. *L. monocytogenes* strains EGD-e (wt), LMSW84 (i*prkA*) and LMS278 (*prkA*ΔC) were grown in BHI broth (containing IPTG where indicated) at 37°C. Average values and standard deviations calculated from technical replicates (n=3) are shown. (C) Increased cephalosporin resistance of a *L. monocytogenes prkA*ΔC mutant. Susceptibility of the same set of strains as in panel B against ceftriaxone was determined using E-tests. Average values and standard deviations were calculated from three independent repetitions. Asterisks mark statistically significant differences (*P*<0.01, *t*-test with Bonferroni-Holm correction). (D) MurA level in an *L. monocytogenes prkA*ΔC mutant. Western blots showing signals specific for MurA (top) in *L. monocytogenes* strains EGD-e (wt), LMJR138 (Δ*clpC*) and LMS278 (*prkA*ΔC). A parallel Western blotting showing DivIVA specific signals was included for control (middle). MurA signals were quantified by densitometry and average values and standard deviations calculated from three independent repetitions are shown (bottom). The asterisk indicates significance level (*P*<0.05, *t*-test with Bonferroni-Holm correction, n=3). (E) Reduced intracellular multiplication of the *prkA*ΔC mutant. The same set of strains as above was used to infect J774 mouse macrophages and bacterial cell numbers were determined right after (0 h p. i.) and 6 hours post infection (6 h p. i.). The experiment was carried out as a triplicate from which average values and standard deviations were calculated. Statistical significance is indicated by asterisks (*P*<0.001, *t*-test with Bonferroni-Holm correction).

### Tn-Seq based identification of genes important during macrophage infection

Next, Tn-Seq was used to identify genes that become essential during infection. For this, J774 mouse macrophages were infected with the Tn insertion library and samples were collected prior to and 24 hours post infection and the number of Tn insertions in each gene before and after infection was determined. All genes with a significant (*P*≤0.05) reduction of Tn-insertions of at least two-fold in two independent experiments were considered as important during intracellular stages. We identified 42 non-essential genes that become essential during infection using this strategy (Tab. 3). First of all, four of the seven virulence genes that form the *Listeria* pathogenicity island LIPI-1 (*prfA*, *plcA*, *hly*, *plcB*) were identified, which demonstrates the suitability of our approach (Fig. 5A, Tab. 3). Remarkably, the two LIPI-1 genes *actA* and *mpl* were not among the identified genes. However, an Δ*actA* mutant was able to replicate in J774 macrophages with wild type kinetic in a separate experiment (Fig. 5B), suggesting that mutants in genuine spreading genes are not counter-selected in our J774 cell culture setup. Besides this, a number of other genes was identified that also had been described previously as essential or important during infection, including genes acting in purine/pyrimidine (*purA*, *purB*, *pyrD*) (17) or menaquinone biosynthesis (*aroE*, *aroF, menB*, *menC*, *menF*, *menI*) (18–20), utilization of branched chain amino acids (*bkdAA*, *bkdAB*) (22), regulation of flagellar motility (*mogR*) (89), daughter cell separation (*divIVA*, *secA2*) (24, 90), lipoate metabolism (*lipL*, *lplA1*) (91, 92) or RNA degradation (*nrnA*) (93) (Tab. 3).

**Figure 5:**
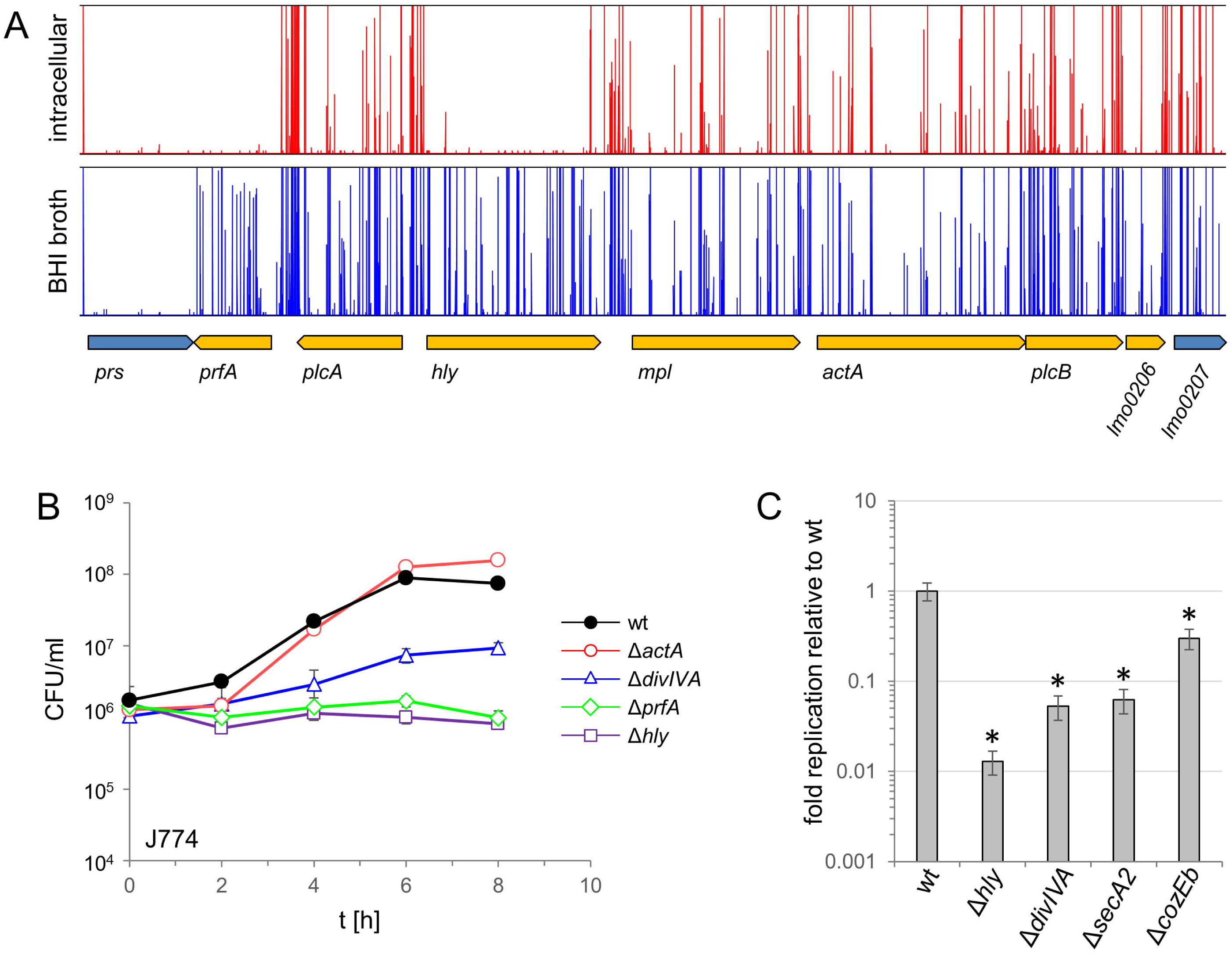
Essentiality of selected virulence genes during intracellular growth in macrophages. (A) Diagrams visualizing the essentiality of LIPI-1 genes during growth under standard growth conditions (BHI broth, blue) and during intracellular growth in J774 mouse macrophages (intracellular, red). Tn-insertions are mapped on the chromosomal region carrying the LIPI-1 genes (*prfA-lmo0206*). Tn-insertions are not found in genes that are essential under a certain condition. The *prs* gene preceding LIPI-1 is essential even under normal growth conditions. (B) Intracellular multiplication of *L. monocytogenes* strains EGD-e (wt), LMS250 (Δ*hly*), LMS251 (Δ*actA*) and BUG2214 (Δ*prfA*) inside J774 mouse macrophages over a time course of 6 h. Strain LMS2 (Δ*divIVA*) was included for comparison. The experiment was performed in triplicate. Average values and standard deviations are shown. (C) Intracellular replication of *L. monocytogenes* strains LMS2 (Δ*divIVA*), LMS81 (Δ*secA2*) and LMJR87 (Δ*cozEb*) mutants in J774 mouse macrophages. Ratio of bacterial titers right after and 6 hours post infection is shown. Wild type replicated 35-fold during this time and was arbitrarily set to 1. Average values and standard deviations were calculated from experiments performed in triplicate. Statistical significance is indicated by asterisks (*P*<0.01, *t*-test with Bonferroni-Holm correction).

**Table 3:**
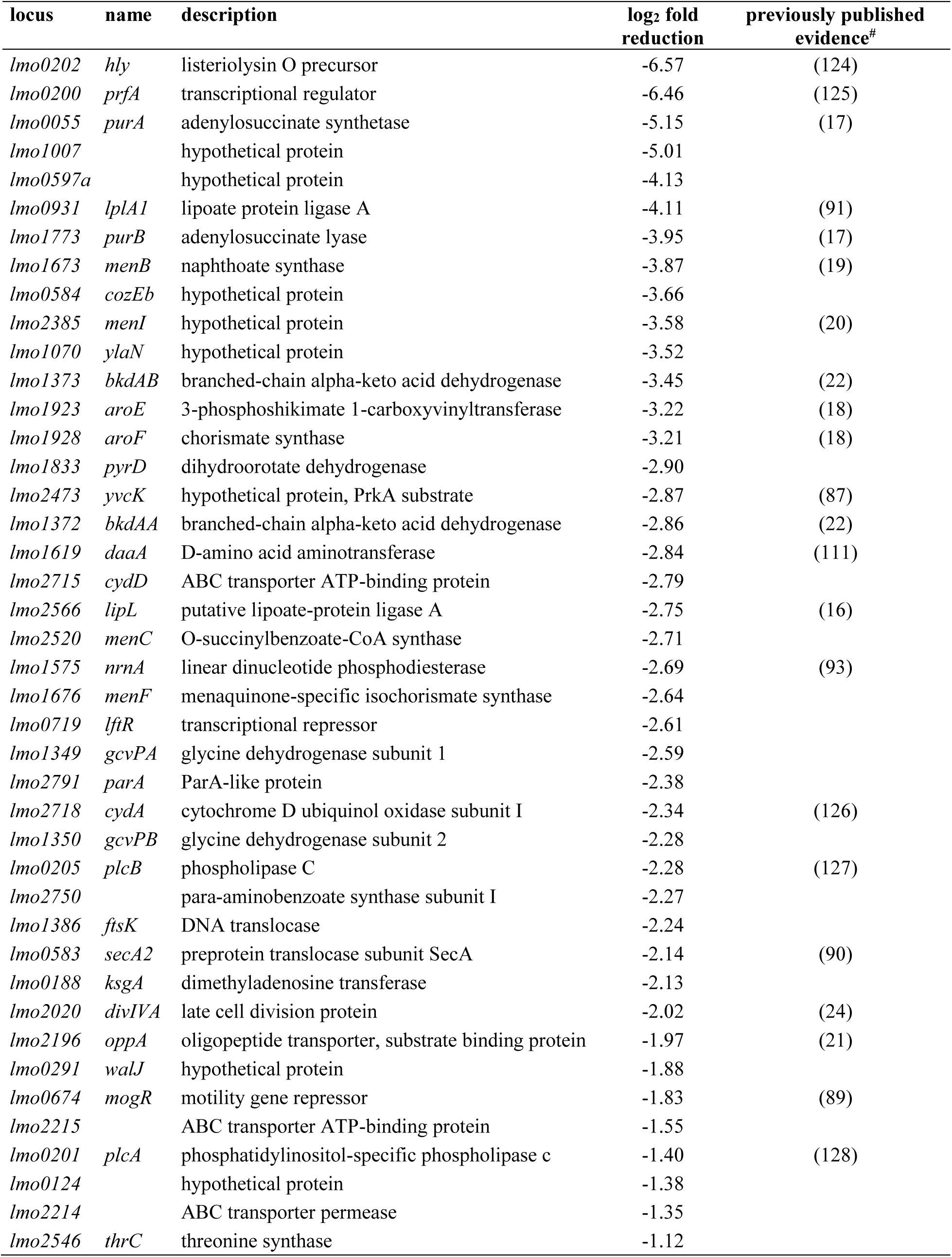
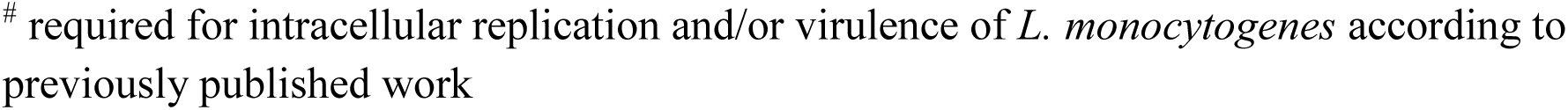
*L. monocytogenes* genes essential during macrophage infection

| locus | name | description | log<sub>2</sub> fold reduction | previously published evidence<sup>#</sup> |
| --- | --- | --- | --- | --- |
| *lmo0202* | *hly* | listeriolysin O precursor | -6.57 | (124) |
| *lmo0200* | *prfA* | transcriptional regulator | -6.46 | (125) |
| *lmo0055* | *purA* | adenylosuccinate synthetase | -5.15 | (17) |
| *lmo1007* |  | hypothetical protein | -5.01 |  |
| *lmo0597a* |  | hypothetical protein | -4.13 |  |
| *lmo0931* | *lplA1* | lipoate protein ligase A | -4.11 | (91) |
| *lmo1773* | *purB* | adenylosuccinate lyase | -3.95 | (17) |
| *lmo1673* | *menB* | naphthoate synthase | -3.87 | (19) |
| *lmo0584* | *cozEb* | hypothetical protein | -3.66 |  |
| *lmo2385* | *menI* | hypothetical protein | -3.58 | (20) |
| *lmo1070* | *ylaN* | hypothetical protein | -3.52 |  |
| *lmo1373* | *bkdAB* | branched-chain alpha-keto acid dehydrogenase | -3.45 | (22) |
| *lmo1923* | *aroE* | 3-phosphoshikimate 1-carboxyvinyltransferase | -3.22 | (18) |
| *lmo1928* | *aroF* | chorismate synthase | -3.21 | (18) |
| *lmo1833* | *pyrD* | dihydrooorotate dehydrogenase | -2.90 |  |
| *lmo2473* | *yvcK* | hypothetical protein, PrkA substrate | -2.87 | (87) |
| *lmo1372* | *bkdAA* | branched-chain alpha-keto acid dehydrogenase | -2.86 | (22) |
| *lmo1619* | *daaA* | D-amino acid aminotransferase | -2.84 | (111) |
| *lmo2715* | *cydD* | ABC transporter ATP-binding protein | -2.79 |  |
| *lmo2566* | *lipL* | putative lipoate-protein ligase A | -2.75 | (16) |
| *lmo2520* | *menC* | O-succinylbenzoate-CoA synthase | -2.71 |  |
| *lmo1575* | *nrnA* | linear dinucleotide phosphodiesterase | -2.69 | (93) |
| *lmo1676* | *menF* | menaquinone-specific isochorismate synthase | -2.64 |  |
| *lmo0719* | *lftR* | transcriptional repressor | -2.61 |  |
| *lmo1349* | *gcvPA* | glycine dehydrogenase subunit 1 | -2.59 |  |
| *lmo2791* | *parA* | ParA-like protein | -2.38 |  |
| *lmo2718* | *cydA* | cytochrome D ubiquinol oxidase subunit I | -2.34 | (126) |
| *lmo1350* | *gcvPB* | glycine dehydrogenase subunit 2 | -2.28 |  |
| *lmo0205* | *plcB* | phospholipase C | -2.28 | (127) |
| *lmo2750* |  | para-aminobenzoate synthase subunit I | -2.27 |  |
| *lmo1386* | *ftsK* | DNA translocase | -2.24 |  |
| *lmo0583* | *secA2* | preprotein translocase subunit SecA | -2.14 | (90) |
| *lmo0188* | *ksgA* | dimethyladenosine transferase | -2.13 |  |
| *lmo2020* | *divIVA* | late cell division protein | -2.02 | (24) |
| *lmo2196* | *oppA* | oligopeptide transporter, substrate binding protein | -1.97 | (21) |
| *lmo0291* | *walJ* | hypothetical protein | -1.88 |  |
| *lmo0674* | *mogR* | motility gene repressor | -1.83 | (89) |
| *lmo2215* |  | ABC transporter ATP-binding protein | -1.55 |  |
| *lmo0201* | *plcA* | phosphatidylinositol-specific phospholipase c | -1.40 | (128) |
| *lmo0124* |  | hypothetical protein | -1.38 |  |
| *lmo2214* |  | ABC transporter permease | -1.35 |  |
| *lmo2546* | *thrC* | threonine synthase | -1.12 |  |

Among the genes so far not linked to intracellular survival were *gcvPA* and *gcvPB*, encoding two components of the glycine cleavage system, the *thrC* gene encoding threonine synthase, as well as the *lmo2214-2215* genes, coding for the components of an ABC transporter of unknown function. Interestingly, several genes with putative functions in PG biosynthesis (*cozEb*, *ylaN*, *yvcK, walJ*) and cell division (*divIVA, ftsK*, *parA*, *secA2*) also showed up in this screen (Tab. 1), out of which some (*divIVA*, *secA2*, *yvcK*) have been identified in previous work (24, 87, 90).

### Novel genes required for intracellular replication

To validate the Tn-Seq results, we created clean deletion mutants in *cozEb*, *walJ*, *ftsK*, *parA*, *gcvPA-gcvPB*, *thrC* and *lmo2214-2215* by allelic replacement. First, replication of the Δ*cozEb* mutant was analysed in J774 macrophages. The Δ*cozEb* mutant generated only 31±16% of the bacterial titre obtained with the wild type after 6 hours of intracellular growth (Fig. 5C), even though no general fitness effect was associated with the *cozEb* deletion in broth culture (Fig. 6A). While this degree of attenuation was clearly less pronounced than that observed with a Δ*hly* mutant (1.2±0.5%), it confirmed the importance of *cozEb* for growth inside macrophages. The *cozEb* gene is located downstream of *secA2* in the chromosome of different *Listeria* species, which together with *divIVA* is crucial for autolysin secretion and virulence (24). The *divIVA* and *secA2* genes were also identified in the screen (Tab. 3), but Δ*divIVA* and Δ*secA2* mutants showed a stronger degree of attenuation (3.7±2.5% and 4.9±3.1% of wild type CFU numbers, respectively) than the Δ*cozEb* mutant (Fig. 5C), suggesting a distinct function for *cozEb*.

**Figure 6:**
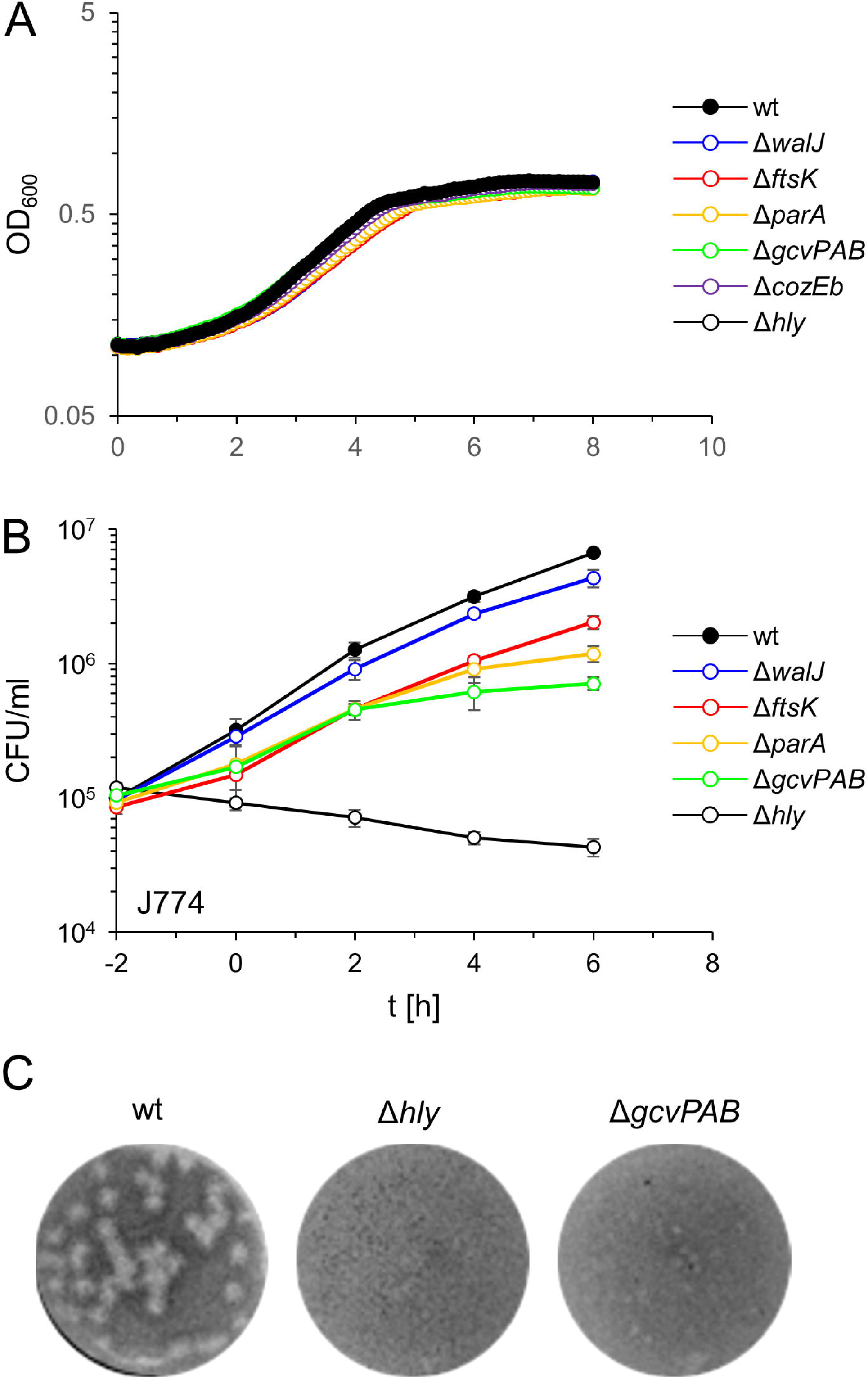
Novel genes contributing to *L. monocytogenes* growth inside macrophages. (A) Growth of *L. monocytogenes* strains EGD-e (wt), LMS250 (Δ*hly*), LMJR87 (Δ*cozEb*), LMS283 (Δ*walJ*), LMS284 (Δ*ftsK*), LMS290 (Δ*parA*) and LMS305 (Δ*gcvPAB*) in BHI broth. (B) Intracellular growth of the same set of strains in J774 mouse macrophages. Average values and standard deviations were calculated from triplicates. (C) Plaque formation assay testing cell-to-cell spread of the Δ*gcvPAB* mutant (strain LMS305) in 3T3 mouse macrophages. *L. monocytogenes* strains EGD-e (wt) and LMS250 (Δ*hly*) were included for control.

Intracellular replication of the remaining mutants was tested in a separate experiment. While mutants lacking the *lmo2214-2215* ABC transporter and *thrC* did not show a delayed intracellular replication (data not shown), the mutant lacking *walJ* showed a modest growth retardation intracellularly (37±27% of wild type CFU numbers after 6 hours, *P*<0.05, Fig. 6B). In contrast, intracellular growth of mutants lacking either *ftsK* (32±24%, *P*<0.01), *parA* (25±19%, *P*<0.005) or *gcvPAB* (20±16%, *P*<0.001) was clearly delayed (Fig. 6B), despite the absence of growth defects in BHI broth (Fig. 6A). Finally, the same strains were tested in plaque formation assays for possible defects in cell-to-cell spread. This revealed a strong defect for the Δ*gcvPAB* mutant (Fig. 6C), while no effects were found for the remaining mutants. Taken together the *cozEb*, *walJ*, *ftsK*, *parA* and *gcvPAB* genes are required for optimal replication in the cytosol of macrophages and *gcvPAB* is additionally essential for cell-to-cell spread.

## DISCUSSION

We here have identified the complement of essential genes that is required to support growth of *Listeria monocytogenes* under standard conditions and during growth in macrophages. With 394 genes, almost 14% of all genes are indispensable for growth in rich medium. Similar numbers of essential genes were discovered by Tn-Seq in other bacteria, such as *Staphylococcus aureus* (420), *Streptococcus pyogenes* (298), *Enterococcus faecalis* (349), *Escherichia coli* (358) or *Pseudomonas aeruginosa* (352 essential genes) (94–98). In *Mycoplasma genitalium*, which contains one of the smallest genomes of self-replicating organisms, 382 out of 482 protein encoding genes are essential (99). We observed a large overlap between the *L. monocytogenes* essential gene set and the 258 essential genes determined by stepwise gene inactivation in *Bacillus subtilis* (100, 101). Moreover, naturally occurring premature stop codons were underrepresented in the *L. monocytogenes* essential genes. Thus, our assignment represents a good estimation of the essential *L. monocytogenes* genome, even though some limitations must be considered. First, a transient incubation at 40°C was required during transposition and this has probably counter-selected heat-sensitive mutants, *e. g.* in *whiA* (this work) or the *gpsB* cell division gene (44). Second, we found several genes as essential by Tn-Seq, even though their inactivation had been reported by others. Heat-sensitivity and different strain backgrounds could be reasons for this. Also, some reported mutants in genes that we consider as essential showed growth defects (*ackA*, *ltaS*, *oppD*, *oppF*, *sipZ*, *sod*, *smpB*, *lmo2487*) (48, 50, 53, 54, 102–104), longer lag phases (*dltA*, *ptsI*) (105) or genetic instability (*spoVG1*) (106). Tn insertions in such genes could also be counter-selected because of their detrimental effects on growth. In other instances, insertional disruptions were reported for genes that we classify as essential, but deletion of these genes (*hprK*, *ptsH*, *pycA*) turned also out to be impossible (52, 107). Probably, these genes are in fact essential, since insertional disruptions are genetically instable and may lead to erroneous conclusions because transient merodiploidy with one wildtype and one mutant allele in the same cell may arise during chromosome replication. Furthermore, suppressor mutations might also have rescued otherwise lethal gene deletions right during strain construction. Thus, it could be interesting to re-examine mutants in genes that we found to be essential but whose deletions were without adverse effects on growth in other studies (*i. e.* the genes shown in Fig. S1) for the possible presence of suppressor mutations and for their heat sensitivity.

Non-surprisingly, ribosomal genes, genes acting in protein biosynthesis and genes with functions in replication, segregation and maintenance of the chromosome constitute the three largest groups of essential genes. Furthermore, stringent patterns of gene essentiality were also observed along central metabolic pathways, first of all glycolysis and the pathways for biosynthesis of the cell envelope. In contrast, the majority of genes for biosynthesis of amino acids and nucleotides was non-essential, which reflects the presence of efficient uptake systems. Most of these genes will become indispensable in minimal medium, where all amino acids (except cysteine for which EGD-e is auxotrophic) need to be synthesized *de novo* using ammonium as the nitrogen source (72). The essentiality of lysine and aspartate biosynthesis genes probably reflect the role of the intermediate/descendant diaminopimelate as a constituent of the peptidoglycan (PG) peptide bridges. PG accounts for a fifth of the weight of a Gram-positive cell (108) and the high number of diaminopimelate molecules needed for this might be difficult to satisfy based on aspartate uptake alone.

The pathways for thiamine, riboflavin, lipoic acid and biotin are incomplete (15, 72) and their few remaining genes are mostly inessential. Consequently, these cofactors must be acquired through essential transporters. In contrast, the genes for biosynthesis of the other cofactors are present, but their essentiality varies. There is a high essentiality ratio in the pathways for coenzyme A, NAD(P)^+^ and tetrahydrofolate (THF) biosynthesis, ruling out the existence of uptake systems, but also disclosing the potential of their enzymes as drug target candidates. The genes for biosynthesis of pyridoxal-phosphate, molybdopterin, porphyrines and ubiquinones, while invariably present, are mostly or partially dispensable. Thus, these compounds or precursors thereof must be taken up via known (porphyrines) or yet to be identified transporters (pyridoxal-phosphate) or could be dispensable during standard growth (ubiquinones and molybdopterin). A haemin transporter is known in *L. monocytogenes* (109) and an ABC transporter for pyridoxal-phosphate was recently discovered in the γ-proteobacterium *Actinobacillus pleuropneumoniae* (110). A homolog of this transporter is not present in *L. monocytogenes*, but one of the many other ABC transporters could execute this function.

Interestingly, only 42 genes become additionally essential when *L. monocytogenes* grows inside macrophages and only half of them act in uptake or biosynthesis of cellular building blocks, suggesting a decent nutrient supply inside the host cell. Among this latter group of genes are such for biosynthesis of certain nucleobases, menaquinones and D-alanine as well as for acquisition of lipoic acid. This is concordant with previous studies (17-19, 91, 111), but it raises the question as to how these compounds contribute to intracellular growth. Scarcity of lipoate and absence of D- alanine explains conditional essentiality of *lplA1*, *lipL* and *daaA* (91, 111). Likewise, a shortage of nucleotide precursors, primarily uracil and xanthine, for the uptake of which many putative transporter genes are present in the genome (112), could explain the intracellular essentiality of the nucleotide biosynthesis genes *purA*, *purB* and *pyrD*. The *aro* and *men* genes are collectively needed for menaquinone biosynthesis and while menaquinones are not required for growth in macrophages (19), one of the intermediates in this pathway, 1,4-dihydroxy-2-naphthoate (DHNA), was shown to be essential for intracellular survival. The specific reason is not clear yet, but DHNA possibly facilitates redox reactions as a cofactor (20).

The glycine cleavage system (GCS) components GcvPA/B were essential intracellularly and also required for cell-to-cell spread. GCS is a lipoate-dependent enzyme system similar to the branched-chain alpha-keto acid dehydrogenase (BKD) enzyme complex, whose genes show the same essentiality pattern. BKD is involved in formation of branched chain fatty acids, which become necessary during infection (22), while GCS transfers a methyl group to THF, the biosynthesis of which is strictly essential. Intracellular essentiality of BKD and GCS would provide an explanation for the importance of lipoate transfer enzymes. GCS might be essential in the host cell since methylated THF is a methyl group donor in other essential reactions (*e. g.* during pantothenate biosynthesis) or because NADH2 generated during glycine degradation is useful as source of energy and electrons.

Our study also highlights the importance of cell division, peptidoglycan biosynthesis and chromosome segregation for extra- and intracellular growth. First of all, several cell division genes are essential (*ezrA*, *ftsA*, *ftsEX*) or heat-sensitive (*whiA*) even though their homologues in *B. subtilis* are not (85, 113–115). Their essential phenotypes may be masked in *B. subtilis* due to a higher degree of gene redundancy, however, essentiality in *L. monocytogenes* may open up new starting points to further investigate the function of these genes in bacterial cell division. Second, several new genes (*cozEb*, *ftsK*, *parA, walJ*) from this functional category have been identified as crucial inside the host cell, even though their function is not understood in all cases. FtsK and ParA are both ATPase needed for chromosome segregation (116), and beyond this, a *L. monocytogenes* Δ*ftsK* mutant was impaired in cell division with a tendency to form chains of unsegregated cells (117). The role of *L. monocytogenes walJ* remains to be studied, but its *B. subtilis* homologue contributes to coordination of cell division with cell wall biosynthesis and chromosome segregation (118). CozEb is a homolog of *S. pneumoniae* CozE, which directs the activity of the penicillin binding protein PBP1a to the MreCD complex to ensure normal PG biosynthesis (119). *S. pneumoniae cozE* is essential, but *L. monocytogenes* encodes two *cozE*-like genes. Possibly, their simultaneous deletion is not tolerated as reported for *S. aureus* (120). Third, our analysis confirms the special role of the two cell division genes *divIVA* and *secA2* for intracellular growth. Deletion of either gene strongly impaired daughter cell separation after septum closure, which leads to formation of long chains of unseparated cells that cannot multiply in macrophages (24, 90). Lastly, we found that the PASTA domains of the essential PrkA kinase can be deleted and that the kinase domain contributes the essential functionality. PrkA regulates PG biosynthesis through phosphorylation of ReoM, a small cytosolic protein that controls proteolytic stability of MurA, the first committed step enzyme in PG biosynthesis, which is essential itself (58, 86, 88). ReoM needs to be phosphorylated to prevent degradation of MurA (58). Deletion of PASTA domains has lowered the activity of StkP, the PrkA homologue of *S. pneumoniae* (121). As MurA can be detected in cells lacking the PASTA domains (Fig. 4D), we assume that the activity of the isolated kinase domain of PrkA is sufficient for ReoM∼P formation. A viable *prkA* mutant with reduced kinase activity will be very useful to further study the role of this protein in *L. monocytogenes* growth and virulence.

## AVAILABILITY

Tn-Seq explorer is a free tool for analysis of Tn-Seq data and available on the GitHub repository: https://github.com/sina-cb/Tn-seqExplorer

The NCBI pathogen detection pipeline collects genome sequences of *L. monocytogenes* isolates worldwide. Data are available at: https://www.ncbi.nlm.nih.gov/pathogens

The cgMLST.org nomenclature server hosts allele profiles of *L. monocytogenes* genes from *L. monocytogenes* isolates of different origin. Data are available at: https://www.cgmlst.org/ncs/schema/690488/

## ACCESSION NUMBERS

Sequencing data was submitted to ENA under project PRJEB51140 as samples ERS10768787 to ERS10768791, each containing the sequencing raw data as well as the used Bam files.

## SUPPLEMENTARY DATA

Supplementary data are available at NAR online.

## ACKNOWLEDGEMENT

We would like to thank Birgitt Hahn for excellent technical assistance and Jeanine Rismondo for help with construction of strain LMJR87. We are also grateful to Kevin S. McIver for kindly sharing plasmid pKRMIT and to Pascale Cossart for sharing the *prfA* mutant.

## REFERENCES

1. Freitag NE, Port GC, Miner MD. *Listeria monocytogenes* - from saprophyte to intracellular pathogen. Nat Rev Microbiol. 2009;7(9):623–8.

2. Grif K, Patscheider G, Dierich MP, Allerberger F. Incidence of fecal carriage of *Listeria monocytogenes* in three healthy volunteers: a one-year prospective stool survey. European journal of clinical microbiology & infectious diseases : official publication of the European Society of Clinical Microbiology. 2003;22(1):16–20.

3. Garcia-Garcera M, Hafner L, Burucoa C, Moura A, Pichon M, Lecuit M. *Listeria monocytogenes* faecal carriage is common and driven by microbiota. bioRxiv. 2021:2021.01.13.426560.

4. Gregory SH, Barczynski LK, Wing EJ. Effector function of hepatocytes and Kupffer cells in the resolution of systemic bacterial infections. J Leukoc Biol. 1992;51(4):421–4.

5. Ebe Y, Hasegawa G, Takatsuka H, Umezu H, Mitsuyama M, Arakawa M, et al. The role of Kupffer cells and regulation of neutrophil migration into the liver by macrophage inflammatory protein-2 in primary listeriosis in mice. Pathol Int. 1999;49(6):519–32.

6. Vazquez-Boland JA, Kuhn M, Berche P, Chakraborty T, Dominguez-Bernal G, Goebel W, et al. *Listeria* pathogenesis and molecular virulence determinants. Clinical microbiology reviews. 2001;14(3):584–640.

7. Cousens LP, Wing EJ. Innate defenses in the liver during *Listeria* infection. Immunol Rev. 2000;174:150–9.

8. Gregory SH, Liu CC. CD8+ T-cell-mediated response to *Listeria monocytogenes* taken up in the liver and replicating within hepatocytes. Immunol Rev. 2000;174:112–22.

9. Lamont RF, Sobel J, Mazaki-Tovi S, Kusanovic JP, Vaisbuch E, Kim SK, et al. Listeriosis in human pregnancy: a systematic review. J Perinat Med. 2011;39(3):227–36.

10. Charlier C, Perrodeau E, Leclercq A, Cazenave B, Pilmis B, Henry B, et al. Clinical features and prognostic factors of listeriosis: the MONALISA national prospective cohort study. Lancet Infect Dis. 2017;17(5):510–9.

11. Scobie A, Kanagarajah S, Harris RJ, Byrne L, Amar C, Grant K, et al. Mortality risk factors for listeriosis - A 10 year review of non-pregnancy associated cases in England 2006-2015. The Journal of infection. 2019;78(3):208–14.

12. Wilking H, Lachmann R, Holzer A, Halbedel S, Flieger A, Stark K. Ongoing high incidence and case fatality of invasive listeriosis in Germany, 2010-2019. Emerging infectious diseases. 2021;accepted.

13. Werber D, Hille K, Frank C, Dehnert M, Altmann D, Müller-Nordhorn J, et al. Years of potential life lost for six major enteric pathogens, Germany, 2004-2008. Epidemiology and infection. 2013;141(5):961–8.

14. Camejo A, Carvalho F, Reis O, Leitao E, Sousa S, Cabanes D. The arsenal of virulence factors deployed by *Listeria monocytogenes* to promote its cell infection cycle. Virulence. 2011;2(5):379–94.

15. Glaser P, Frangeul L, Buchrieser C, Rusniok C, Amend A, Baquero F, et al. Comparative genomics of *Listeria* species. Science. 2001;294(5543):849–52.

16. Schauer K, Geginat G, Liang C, Goebel W, Dandekar T, Fuchs TM. Deciphering the intracellular metabolism of *Listeria monocytogene*s by mutant screening and modelling. BMC Genomics. 2010;11:573.

17. Faith NG, Kim JW, Azizoglu R, Kathariou S, Czuprynski C. Purine biosynthesis mutants (purA and purB) of serotype 4b *Listeria monocytogenes* are severely attenuated for systemic infection in intragastrically inoculated A/J Mice. Foodborne Pathog Dis. 2012;9(5):480–6.

18. Stritzker J, Janda J, Schoen C, Taupp M, Pilgrim S, Gentschev I, et al. Growth, virulence, and immunogenicity of *Listeria monocytogenes* aro mutants. Infect Immun. 2004;72(10):5622–9.

19. Chen GY, McDougal CE, D’Antonio MA, Portman JL, Sauer JD. A Genetic Screen Reveals that Synthesis of 1,4-Dihydroxy-2-Naphthoate (DHNA), but Not Full-Length Menaquinone, Is Required for *Listeria monocytogenes* Cytosolic Survival. mBio. 2017;8(2).

20. Smith HB, Li TL, Liao MK, Chen GY, Guo Z, Sauer JD. *Listeria monocytogene*s MenI Encodes a DHNA-CoA Thioesterase Necessary for Menaquinone Biosynthesis, Cytosolic Survival, and Virulence. Infect Immun. 2021;89(5).

21. Borezee E, Pellegrini E, Berche P. OppA of *Listeria monocytogenes*, an oligopeptide-binding protein required for bacterial growth at low temperature and involved in intracellular survival. Infect Immun. 2000;68(12):7069–77.

22. Sun Y, O’Riordan MX. Branched-chain fatty acids promote *Listeria monocytogenes* intracellular infection and virulence. Infect Immun. 2010;78(11):4667–73.

23. Lenz LL, Portnoy DA. Identification of a second *Listeria secA* gene associated with protein secretion and the rough phenotype. Mol Microbiol. 2002;45(4):1043–56.

24. Halbedel S, Hahn B, Daniel RA, Flieger A. DivIVA affects secretion of virulence-related autolysins in *Listeria monocytogenes*. Mol Microbiol. 2012;83(4):821–39.

25. Joseph B, Przybilla K, Stuhler C, Schauer K, Slaghuis J, Fuchs TM, et al. Identification of *Listeria monocytogenes* genes contributing to intracellular replication by expression profiling and mutant screening. J Bacteriol. 2006;188(2):556–68.

26. van Opijnen T, Bodi KL, Camilli A. Tn-seq: high-throughput parallel sequencing for fitness and genetic interaction studies in microorganisms. Nature methods. 2009;6(10):767–72.

27. Sambrook J, Fritsch EF, Maniatis T. Molecular cloning : a laboratory manual. 2nd ed. Cold Spring Harbor, N.Y.: Cold Spring Harbor Laboratory Press; 1989. 3 v. p.

28. Monk IR, Gahan CG, Hill C. Tools for functional postgenomic analysis of *Listeria monocytogenes*. Appl Environ Microbiol. 2008;74(13):3921–34.

29. Martin M. Cutadapt removes adapter sequences from high-throughput sequencing reads. 2011. 2011;17(1):3.

30. Toledo-Arana A, Dussurget O, Nikitas G, Sesto N, Guet-Revillet H, Balestrino D, et al. The *Listeria* transcriptional landscape from saprophytism to virulence. Nature. 2009;459(7249):950–6.

31. Langmead B, Trapnell C, Pop M, Salzberg SL. Ultrafast and memory-efficient alignment of short DNA sequences to the human genome. Genome Biol. 2009;10(3):R25.

32. Solaimanpour S, Sarmiento F, Mrazek J. Tn-seq explorer: a tool for analysis of high-throughput sequencing data of transposon mutant libraries. PLoS One. 2015;10(5):e0126070.

33. DeJesus MA, Ambadipudi C, Baker R, Sassetti C, Ioerger TR. TRANSIT--A Software Tool for Himar1 TnSeq Analysis. PLoS Comput Biol. 2015;11(10):e1004401.

34. Carver T, Harris SR, Berriman M, Parkhill J, McQuillan JA. Artemis: an integrated platform for visualization and analysis of high-throughput sequence-based experimental data. Bioinformatics. 2012;28(4):464–9.

35. Ruppitsch W, Pietzka A, Prior K, Bletz S, Fernandez HL, Allerberger F, et al. Defining and Evaluating a Core Genome Multilocus Sequence Typing Scheme for Whole-Genome Sequence-Based Typing of *Listeria monocytogenes*. J Clin Microbiol. 2015;53(9):2869–76.

36. van den Ent F, Löwe J. RF cloning: a restriction-free method for inserting target genes into plasmids. J Biochem Biophys Methods. 2006;67(1):67–74.

37. Arnaud M, Chastanet A, Debarbouille M. New vector for efficient allelic replacement in naturally nontransformable, low-GC-content, gram-positive bacteria. Appl Environ Microbiol. 2004;70(11):6887–91.

38. Marston AL, Thomaides HB, Edwards DH, Sharpe ME, Errington J. Polar localization of the MinD protein of *Bacillus subtilis* and its role in selection of the mid-cell division site. Genes Dev. 1998;12(21):3419–30.

39. Kock H, Gerth U, Hecker M. MurAA, catalysing the first committed step in peptidoglycan biosynthesis, is a target of Clp-dependent proteolysis in *Bacillus subtilis*. Mol Microbiol. 2004;51(4):1087–102.

40. Halbedel S, Reiss S, Hahn B, Albrecht D, Mannala GK, Chakraborty T, et al. A systematic proteomic analysis of *Listeria monocytogenes* house-keeping protein secretion systems. Molecular & cellular proteomics : MCP. 2014;13(11):3063–81.

41. Langridge GC, Phan MD, Turner DJ, Perkins TT, Parts L, Haase J, et al. Simultaneous assay of every *Salmonella* Typhi gene using one million transposon mutants. Genome Res. 2009;19(12):2308–16.

42. Hauf S, Herrmann J, Miethke M, Gibhardt J, Commichau FM, Müller R, et al. Aurantimycin resistance genes contribute to survival of *Listeria monocytogenes* during life in the environment. Mol Microbiol. 2019;111(4):1009–24.

43. Zhu B, Stülke J. SubtiWiki in 2018: from genes and proteins to functional network annotation of the model organism *Bacillus subtilis*. Nucleic Acids Res. 2018;46(D1):D743–D8.

44. Rismondo J, Cleverley RM, Lane HV, Grosshennig S, Steglich A, Möller L, et al. Structure of the bacterial cell division determinant GpsB and its interaction with penicillin-binding proteins. Mol Microbiol. 2016;99(5):978–98.

45. Gaillot O, Pellegrini E, Bregenholt S, Nair S, Berche P. The ClpP serine protease is essential for the intracellular parasitism and virulence of *Listeria monocytogenes*. Mol Microbiol. 2000;35(6):1286–94.

46. Hanawa T, Fukuda M, Kawakami H, Hirano H, Kamiya S, Yamamoto T. The *Listeria monocytogenes* DnaK chaperone is required for stress tolerance and efficient phagocytosis with macrophages. Cell Stress Chaperones. 1999;4(2):118–28.

47. Markkula A, Lindstrom M, Johansson P, Bjorkroth J, Korkeala H. Roles of four putative DEAD-box RNA helicase genes in growth of *Listeria monocytogenes* EGD-e under heat, pH, osmotic, ethanol, and oxidative stress conditions. Appl Environ Microbiol. 2012;78(19):6875–82.

48. Gueriri I, Bay S, Dubrac S, Cyncynatus C, Msadek T. The Pta-AckA pathway controlling acetyl phosphate levels and the phosphorylation state of the DegU orphan response regulator both play a role in regulating *Listeria monocytogenes* motility and chemotaxis. Mol Microbiol. 2008;70(6):1342–57.

49. Webb AJ, Karatsa-Dodgson M, Gründling A. Two-enzyme systems for glycolipid and polyglycerolphosphate lipoteichoic acid synthesis in *Listeria monocytogenes*. Mol Microbiol. 2009;74(2):299–314.

50. Krypotou E, Scortti M, Grundstrom C, Oelker M, Luisi BF, Sauer-Eriksson AE, et al. Control of Bacterial Virulence through the Peptide Signature of the Habitat. Cell reports. 2019;26(7):1815–27 e5.

51. Ake FM, Joyet P, Deutscher J, Milohanic E. Mutational analysis of glucose transport regulation and glucose-mediated virulence gene repression in *Listeria monocytogenes*. Mol Microbiol. 2011;81(1):274–93.

52. Schar J, Stoll R, Schauer K, Loeffler DI, Eylert E, Joseph B, et al. Pyruvate carboxylase plays a crucial role in carbon metabolism of extra- and intracellularly replicating *Listeria monocytogenes*. J Bacteriol. 2010;192(7):1774–84.

53. Bonnemain C, Raynaud C, Reglier-Poupet H, Dubail I, Frehel C, Lety MA, et al. Differential roles of multiple signal peptidases in the virulence of *Listeria monocytogenes*. Mol Microbiol. 2004;51(5):1251–66.

54. Archambaud C, Nahori MA, Pizarro-Cerda J, Cossart P, Dussurget O. Control of *Listeria* superoxide dismutase by phosphorylation. J Biol Chem. 2006;281(42):31812–22.

55. Considine KM, Sleator RD, Kelly AL, Fitzgerald GF, Hill C. Identification and characterization of an essential gene in *Listeria monocytogenes* using an inducible gene expression system. Bioengineered bugs. 2011;2(3):150–9.

56. Woodward JJ, Iavarone AT, Portnoy DA. c-di-AMP secreted by intracellular *Listeria monocytogenes* activates a host type I interferon response. Science. 2010;328(5986):1703–5.

57. Rismondo J, Möller L, Aldridge C, Gray J, Vollmer W, Halbedel S. Discrete and overlapping functions of peptidoglycan synthases in growth, cell division and virulence of *Listeria monocytogenes*. Mol Microbiol. 2015;95(2):332–51.

58. Wamp S, Rutter ZJ, Rismondo J, Jennings CE, Möller L, Lewis RJ, et al. PrkA controls peptidoglycan biosynthesis through the essential phosphorylation of ReoM. eLife. 2020;9.

59. Bennett HJ, Pearce DM, Glenn S, Taylor CM, Kuhn M, Sonenshein AL, et al. Characterization of *relA* and *codY* mutants of *Listeria monocytogenes*: identification of the CodY regulon and its role in virulence. Mol Microbiol. 2007;63(5):1453–67.

60. van der Veen S, Abee T. Contribution of *Listeria monocytogenes* RecA to acid and bile survival and invasion of human intestinal Caco-2 cells. Int J Med Microbiol. 2011;301(4):334–40.

61. Burg-Golani T, Pozniak Y, Rabinovich L, Sigal N, Nir Paz R, Herskovits AA. Membrane chaperone SecDF plays a role in the secretion of *Listeria monocytogenes* major virulence factors. J Bacteriol. 2013;195(23):5262–72.

62. McLaughlin HP, Xiao Q, Rea RB, Pi H, Casey PG, Darby T, et al. A putative P-type ATPase required for virulence and resistance to haem toxicity in *Listeria monocytogenes*. PLoS One. 2012;7(2):e30928.

63. Markkula A, Mattila M, Lindstrom M, Korkeala H. Genes encoding putative DEAD-box RNA helicases in *Listeria monocytogenes* EGD-e are needed for growth and motility at 3 degrees C. Environ Microbiol. 2012;14(8):2223–32.

64. NCBI. The NCBI Pathogen Detection Project [Internet] Bethesda (MD): National Library of Medicine (US), National Center for Biotechnology Information; 2020 [Available from: https://www.ncbi.nlm.nih.gov/pathogens/.

65. Tolentino R, Chastain C, Burnell J. Identification of the amino acid involved in the regulation of bacterial pyruvate, orthophosphate dikinase and phosphoenolpyruvate synthetase. Advances in Biological Chemistry. 2013;3(3A):12–21.

66. Cabrera-Valladares N, Martinez LM, Flores N, Hernandez-Chavez G, Martinez A, Bolivar F, et al. Physiologic consequences of glucose transport and phosphoenolpyruvate node modifications in *Bacillus subtilis* 168. J Mol Microbiol Biotechnol. 2012;22(3):177–97.

67. Pine L, Malcolm GB, Brooks JB, Daneshvar MI. Physiological studies on the growth and utilization of sugars by *Listeria* species. Can J Microbiol. 1989;35(2):245–54.

68. Romick TL, Fleming HP, McFeeters RF. Aerobic and anaerobic metabolism of *Listeria monocytogenes* in defined glucose medium. Appl Environ Microbiol. 1996;62(1):304–7.

69. Trivett TL, Meyer EA. Citrate cycle and related metabolism of *Listeria monocytogenes*. J Bacteriol. 1971;107(3):770–9.

70. Kentache T, Milohanic E, Cao TN, Mokhtari A, Ake FM, Ma Pham QM, et al. Transport and Catabolism of Pentitols by *Listeria monocytogenes*. J Mol Microbiol Biotechnol. 2016;26(6):369–80.

71. Yao J, Ericson ME, Frank MW, Rock CO. Enoyl-Acyl Carrier Protein Reductase I (FabI) Is Essential for the Intracellular Growth of *Listeria monocytogenes*. Infect Immun. 2016;84(12):3597–607.

72. Tsai HN, Hodgson DA. Development of a synthetic minimal medium for *Listeria monocytogenes*. Appl Environ Microbiol. 2003;69(11):6943–5.

73. Kutzner E, Kern T, Felsl A, Eisenreich W, Fuchs TM. Isotopologue profiling of the listerial N-metabolism. Mol Microbiol. 2016;100(2):315–27.

74. Premaratne RJ, Lin WJ, Johnson EA. Development of an improved chemically defined minimal medium for *Listeria monocytogenes*. Appl Environ Microbiol. 1991;57(10):3046–8.

75. Matern A, Pedrolli D, Grosshennig S, Johansson J, Mack M. Uptake and Metabolism of Antibiotics Roseoflavin and 8-Demethyl-8-Aminoriboflavin in Riboflavin-Auxotrophic *Listeria monocytogenes*. J Bacteriol. 2016;198(23):3233–43.

76. Schauer K, Stolz J, Scherer S, Fuchs TM. Both thiamine uptake and biosynthesis of thiamine precursors are required for intracellular replication of *Listeria monocytogenes*. J Bacteriol. 2009;191(7):2218–27.

77. Dowd GC, Joyce SA, Hill C, Gahan CG. Investigation of the mechanisms by which *Listeria monocytogenes* grows in porcine gallbladder bile. Infect Immun. 2011;79(1):369–79.

78. Belitsky BR. Role of PdxR in the activation of vitamin B6 biosynthesis in *Listeria monocytogenes*. Mol Microbiol. 2014;92(5):1113–28.

79. Richts B, Rosenberg J, Commichau FM. A Survey of Pyridoxal 5’-Phosphate-Dependent Proteins in the Gram-Positive Model Bacterium *Bacillus subtilis*. Front Mol Biosci. 2019;6:32.

80. Klebba PE, Charbit A, Xiao Q, Jiang X, Newton SM. Mechanisms of iron and haem transport by *Listeria monocytogenes*. Molecular membrane biology. 2012;29(3-4):69–86.

81. Rismondo J, Halbedel S, Gründling A. Cell Shape and Antibiotic Resistance Are Maintained by the Activity of Multiple FtsW and RodA Enzymes in *Listeria monocytogenes*. mBio. 2019;10(4).

82. Eugster MR, Loessner MJ. Wall teichoic acids restrict access of bacteriophage endolysin Ply118, Ply511, and PlyP40 cell wall binding domains to the *Listeria monocytogenes* peptidoglycan. J Bacteriol. 2012;194(23):6498–506.

83. Carvalho F, Atilano ML, Pombinho R, Covas G, Gallo RL, Filipe SR, et al. L-Rhamnosylation of Listeria monocytogenes wall teichoic acids promotes resistance to antimicrobial peptides by delaying interaction with the membrane. PLoS Pathog. 2015;11(5):e1004919.

84. Rismondo J, Percy MG, Gründling A. Discovery of genes required for lipoteichoic acid glycosylation predicts two distinct mechanism for wall teichoic acid glycosylation. J Biol Chem. 2018.

85. Surdova K, Gamba P, Claessen D, Siersma T, Jonker MJ, Errington J, et al. The conserved DNA-binding protein WhiA is involved in cell division in *Bacillus subtilis*. J Bacteriol. 2013;195(24):5450–60.

86. Wamp S, Rothe P, Holland G, Halbedel S. MurA escape mutations uncouple peptidoglycan biosynthesis from PrkA signaling. bioRxiv. 2021:2021.09.09.459578.

87. Pensinger DA, Boldon KM, Chen GY, Vincent WJ, Sherman K, Xiong M, et al. The *Listeria monocytogenes* PASTA Kinase PrkA and its substrate YvcK are required for cell wall homeostasis, metabolism, and virulence. PLoS Pathog. 2016;12(11):e1006001.

88. Kelliher JL, Grunenwald CM, Abrahams RR, Daanen ME, Lew CI, Rose WE, et al. PASTA kinase-dependent control of peptidoglycan synthesis via ReoM is required for cell wall stress responses, cytosolic survival, and virulence in *Listeria monocytogenes*. PLoS Pathog. 2021;17(10):e1009881.

89. Gründling A, Burrack LS, Bouwer HG, Higgins DE. *Listeria monocytogenes* regulates flagellar motility gene expression through MogR, a transcriptional repressor required for virulence. Proc Natl Acad Sci U S A. 2004;101(33):12318–23.

90. Lenz LL, Mohammadi S, Geissler A, Portnoy DA. SecA2-dependent secretion of autolytic enzymes promotes *Listeria monocytogenes* pathogenesis. Proc Natl Acad Sci U S A. 2003;100(21):12432–7.

91. O’Riordan M, Moors MA, Portnoy DA. *Listeria* intracellular growth and virulence require host- derived lipoic acid. Science. 2003;302(5644):462–4.

92. Dowd GC, Bahey-El-Din M, Casey PG, Joyce SA, Hill C, Gahan CG. *Listeria monocytogenes* mutants defective in gallbladder replication represent safety-enhanced vaccine delivery platforms. Human vaccines & immunotherapeutics. 2016;12(8):2059–63.

93. Gall AR, Hsueh BY, Siletti C, Waters CM, Huynh TN. NrnA is a linear dinucleotide phosphodiesterase with limited function in cyclic dinucleotide metabolism in *Listeria monocytogenes*. J Bacteriol. 2021:JB0020621.

94. Valentino MD, Foulston L, Sadaka A, Kos VN, Villet RA, Santa Maria J, Jr., et al. Genes contributing to *Staphylococcus aureus* fitness in abscess- and infection-related ecologies. mBio. 2014;5(5):e01729–14.

95. Le Breton Y, Belew AT, Valdes KM, Islam E, Curry P, Tettelin H, et al. Essential Genes in the Core Genome of the Human Pathogen *Streptococcus pyogenes*. Scientific reports. 2015;5:9838.

96. Gilmore MS, Salamzade R, Selleck E, Bryan N, Mello SS, Manson AL, et al. Genes Contributing to the Unique Biology and Intrinsic Antibiotic Resistance of *Enterococcus faecalis*. mBio. 2020;11(6).

97. Goodall ECA, Robinson A, Johnston IG, Jabbari S, Turner KA, Cunningham AF, et al. The Essential Genome of Escherichia coli K-12. mBio. 2018;9(1).

98. Lee SA, Gallagher LA, Thongdee M, Staudinger BJ, Lippman S, Singh PK, et al. General and condition-specific essential functions of *Pseudomonas aeruginosa*. Proc Natl Acad Sci U S A. 2015;112(16):5189–94.

99. Glass JI, Assad-Garcia N, Alperovich N, Yooseph S, Lewis MR, Maruf M, et al. Essential genes of a minimal bacterium. Proc Natl Acad Sci U S A. 2006;103(2):425–30.

100. Kobayashi K, Ehrlich SD, Albertini A, Amati G, Andersen KK, Arnaud M, et al. Essential *Bacillus subtilis* genes. Proc Natl Acad Sci U S A. 2003;100(8):4678–83.

101. Commichau FM, Pietack N, Stülke J. Essential genes in *Bacillus subtilis*: a re-evaluation after ten years. Molecular bioSystems. 2013;9(6):1068–75.

102. Webb AJ, Karatsa-Dodgson M, Grundling A. Two-enzyme systems for glycolipid and polyglycerolphosphate lipoteichoic acid synthesis in *Listeria monocytogenes*. Mol Microbiol. 2009;74(2):299–314.

103. Mraheil MA, Frantz R, Teubner L, Wendt H, Linne U, Wingerath J, et al. Requirement of the RNA-binding protein SmpB during intracellular growth of *Listeria monocytogenes*. Int J Med Microbiol. 2017;307(3):166–73.

104. Gravesen A, Kallipolitis B, Holmstrom K, Hoiby PE, Ramnath M, Knochel S. *pbp2229*-mediated nisin resistance mechanism in *Listeria monocytogenes* confers cross-protection to class IIa bacteriocins and affects virulence gene expression. Appl Environ Microbiol. 2004;70(3):1669–79.

105. Abachin E, Poyart C, Pellegrini E, Milohanic E, Fiedler F, Berche P, et al. Formation of D-alanyl-lipoteichoic acid is required for adhesion and virulence of *Listeria monocytogenes*. Mol Microbiol. 2002;43(1):1–14.

106. Burke TP, Portnoy DA. SpoVG Is a Conserved RNA-Binding Protein That Regulates *Listeria monocytogenes* Lysozyme Resistance, Virulence, and Swarming Motility. mBio. 2016;7(2):e00240.

107. Mertins S, Joseph B, Goetz M, Ecke R, Seidel G, Sprehe M, et al. Interference of components of the phosphoenolpyruvate phosphotransferase system with the central virulence gene regulator PrfA of *Listeria monocytogenes*. J Bacteriol. 2007;189(2):473–90.

108. Reith J, Mayer C. Peptidoglycan turnover and recycling in Gram-positive bacteria. Appl Microbiol Biotechnol. 2011;92(1):1–11.

109. Xiao Q, Jiang X, Moore KJ, Shao Y, Pi H, Dubail I, et al. Sortase independent and dependent systems for acquisition of haem and haemoglobin in *Listeria monocytogenes*. Mol Microbiol. 2011;80(6):1581–97.

110. Pan C, Zimmer A, Shah M, Huynh MS, Lai CC, Sit B, et al. *Actinobacillu*s utilizes a binding protein-dependent ABC transporter to acquire the active form of vitamin B6. J Biol Chem. 2021;297(3):101046.

111. Thompson RJ, Bouwer HG, Portnoy DA, Frankel FR. Pathogenicity and immunogenicity of a *Listeria monocytogenes* strain that requires D-alanine for growth. Infect Immun. 1998;66(8):3552–61.

112. Elbourne LD, Tetu SG, Hassan KA, Paulsen IT. TransportDB 2.0: a database for exploring membrane transporters in sequenced genomes from all domains of life. Nucleic Acids Res. 2017;45(D1):D320–D4.

113. Levin PA, Kurtser IG, Grossman AD. Identification and characterization of a negative regulator of FtsZ ring formation in *Bacillus subtilis*. Proc Natl Acad Sci U S A. 1999;96(17):9642–7.

114. Beall B, Lutkenhaus J. Nucleotide sequence and insertional inactivation of a *Bacillus subtilis* gene that affects cell division, sporulation, and temperature sensitivity. J Bacteriol. 1989;171(12):6821–34.

115. Garti-Levi S, Hazan R, Kain J, Fujita M, Ben-Yehuda S. The FtsEX ABC transporter directs cellular differentiation in *Bacillus subtilis*. Mol Microbiol. 2008;69(4):1018–28.

116. Bouet JY, Stouf M, Lebailly E, Cornet F. Mechanisms for chromosome segregation. Curr Opin Microbiol. 2014;22:60–5.

117. Chang Y, Gu W, Zhang F, McLandsborough L. Disruption of *lmo1386*, a putative DNA translocase gene, affects biofilm formation of *Listeria monocytogenes* on abiotic surfaces. Int J Food Microbiol. 2013;161(3):158–63.

118. Biller SJ, Wayne KJ, Winkler ME, Burkholder WF. The putative hydrolase YycJ (WalJ) affects the coordination of cell division with DNA replication in *Bacillus subtilis* and may play a conserved role in cell wall metabolism. J Bacteriol. 2011;193(4):896–908.

119. Fenton AK, El Mortaji L, Lau DT, Rudner DZ, Bernhardt TG. CozE is a member of the MreCD complex that directs cell elongation in *Streptococcus pneumoniae*. Nat Microbiol. 2016;2:16237.

120. Stamsas GA, Myrbraten IS, Straume D, Salehian Z, Veening JW, Havarstein LS, et al. CozEa and CozEb play overlapping and essential roles in controlling cell division in *Staphylococcus aureus*. Mol Microbiol. 2018;109(5):615–32.

121. Zucchini L, Mercy C, Garcia PS, Cluzel C, Gueguen-Chaignon V, Galisson F, et al. PASTA repeats of the protein kinase StkP interconnect cell constriction and separation of *Streptococcus pneumoniae*. Nat Microbiol. 2018;3(2):197–209.

122. Mandin P, Repoila F, Vergassola M, Geissmann T, Cossart P. Identification of new noncoding RNAs in *Listeria monocytogenes* and prediction of mRNA targets. Nucleic Acids Res. 2007;35(3):962–74.

123. Rismondo J, Bender JK, Halbedel S. Suppressor Mutations Linking gpsB with the First Committed Step of Peptidoglycan Biosynthesis in *Listeria monocytogenes*. J Bacteriol. 2017;199(1).

124. Portnoy DA, Jacks PS, Hinrichs DJ. Role of hemolysin for the intracellular growth of *Listeria monocytogenes*. The Journal of experimental medicine. 1988;167(4):1459–71.

125. Chakraborty T, Leimeister-Wächter M, Domann E, Hartl M, Goebel W, Nichterlein T, et al. Coordinate regulation of virulence genes in *Listeria monocytogenes* requires the product of the *prfA* gene. J Bacteriol. 1992;174(2):568–74.

126. Corbett D, Goldrick M, Fernandes VE, Davidge K, Poole RK, Andrew PW, et al. *Listeria monocytogenes* Has Both Cytochrome bd-Type and Cytochrome aa 3-Type Terminal Oxidases, Which Allow Growth at Different Oxygen Levels, and Both Are Important in Infection. Infect Immun. 2017;85(11).

127. Vazquez-Boland JA, Kocks C, Dramsi S, Ohayon H, Geoffroy C, Mengaud J, et al. Nucleotide sequence of the lecithinase operon of *Listeria monocytogenes* and possible role of lecithinase in cell-to-cell spread. Infect Immun. 1992;60(1):219–30.

128. Camilli A, Goldfine H, Portnoy DA. *Listeria monocytogenes* mutants lacking phosphatidylinositol- specific phospholipase C are avirulent. The Journal of experimental medicine. 1991;173(3):751–4.

