## Supplemental Figures S1-S5 for "*Listeria monocytogenes* gene essentiality under laboratory conditions and during macrophage infection"

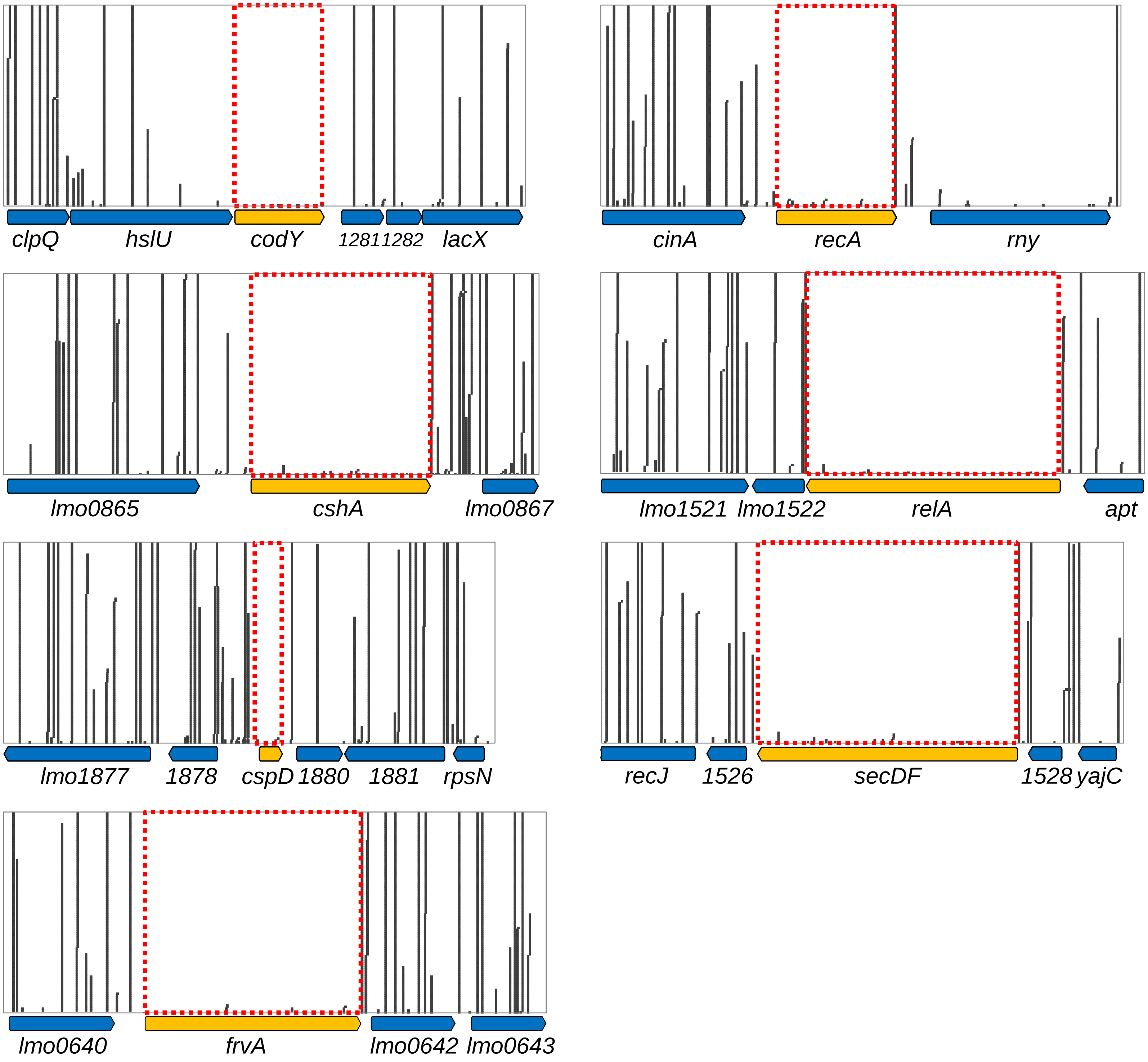


**Fig. S1:** Absence of Tn insertions in genes that could be deleted in previous studies.
Distribution of Tn insertions at the *codY*, *cshA*, *cspD*, *frvA*, *recA*, *relA* and *secDF* loci of Tn mutagenized *L. monocytogenes* EDG-e, which resulted in the categorization as essential gene during standard growth conditions in this study.

Fischer *et al.* Fig. S1

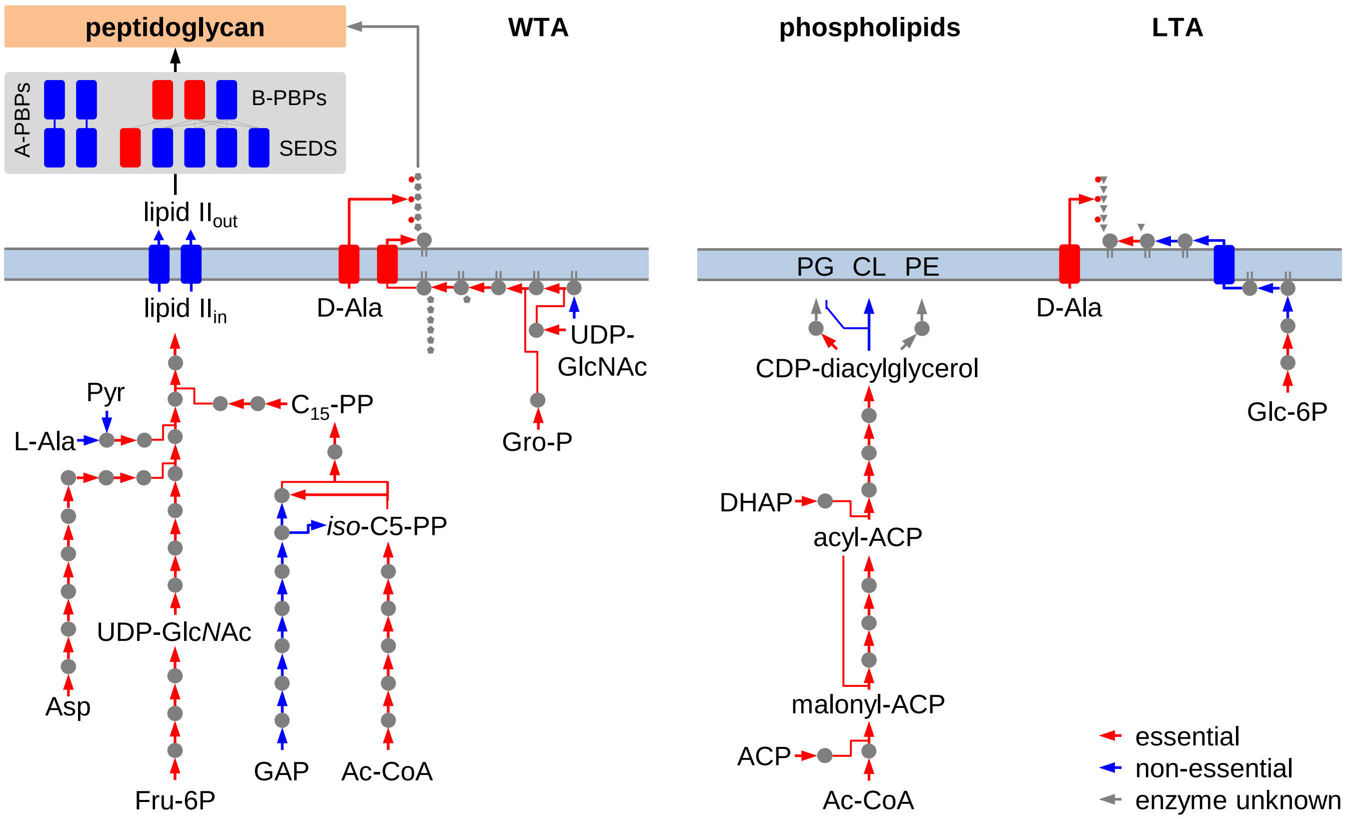


**Fig. S2:** Gene essentiality in biosynthetic pathways for envelope biosynthesis.
Schemes illustrating the *L. monocytogenes* pathways for biosynthesis of peptidoglycan and wall teichoic acids (WTA, left), phospholipids (middle) and lipoteichoic acids (LTA, right) according to the KEGG database (https://www.genome.jp/kegg-bin/show_organism?org=T00066). Reactions are colored according to the essentiality of their corresponding genes. Pathways for LTA and WTA glycosylation were not included.

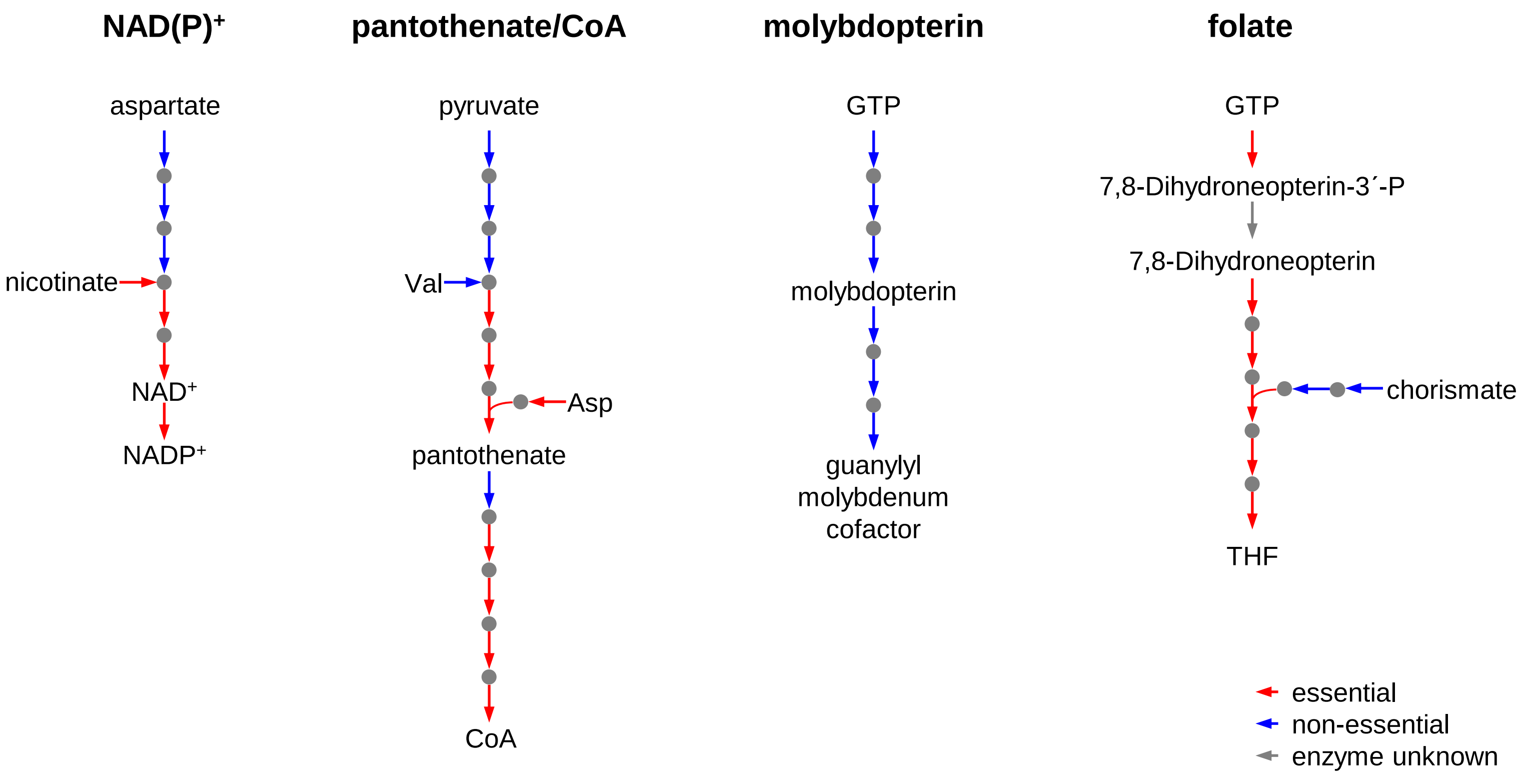


**Fig. S3:** Gene essentiality in biosynthetic pathways for selected co-factors.
Schemes illustrating the anabolic pathways for biosynthesis of NAD(P)$^+$, coenzyme A, molybdopterin and folate according to the KEGG database (https://www.genome.jp/kegg-bin/show_organism?org=T00066). Reactions are colored according to the essentiality of their corresponding genes.

Fischer *et al.* Fig. S3

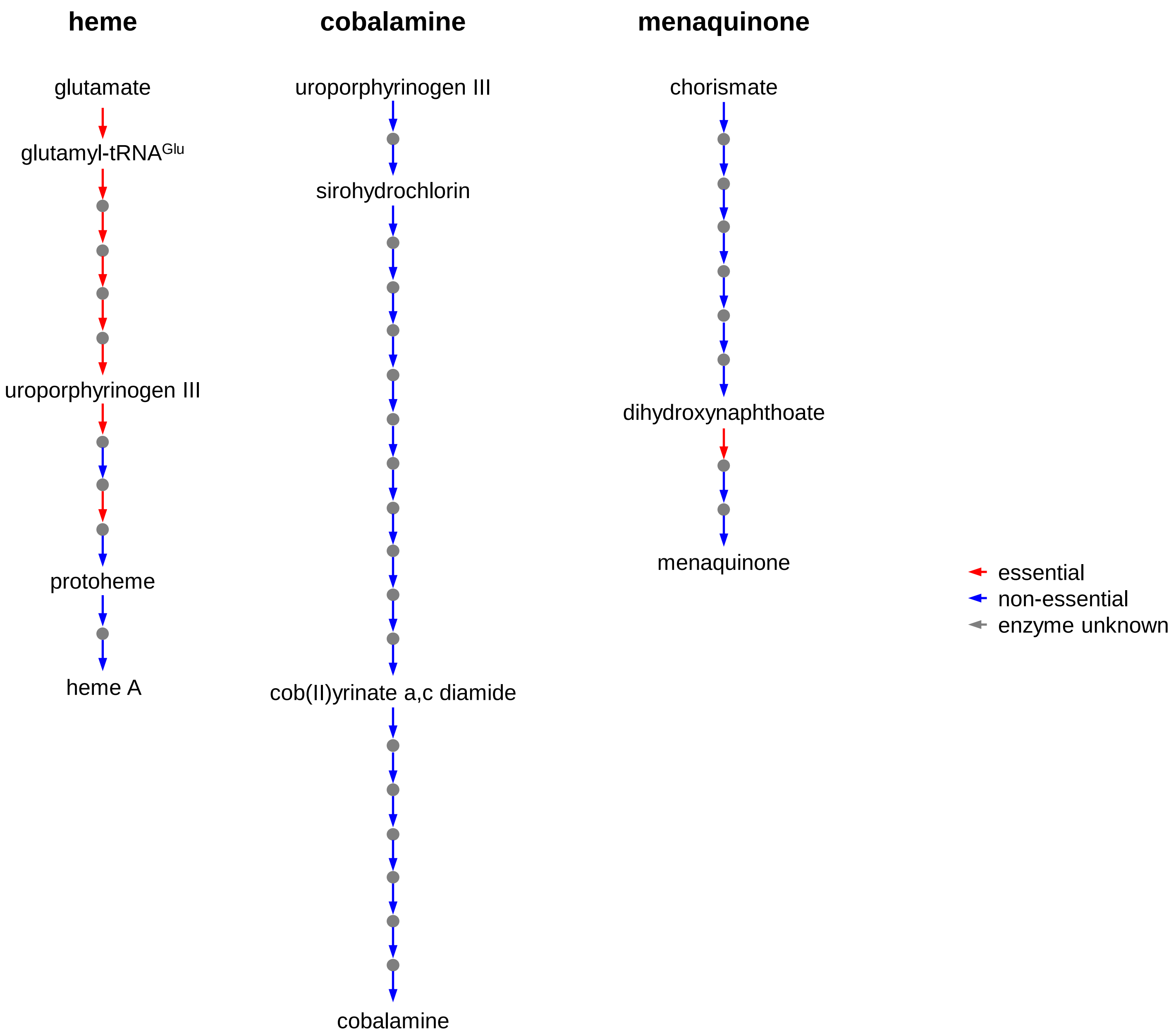


**Fig. S4:** Gene essentiality in biosynthetis for heme, cobalamine and menaquinone.
Schemes illustrating the anabolic pathways for heme, cobalamine and menaquinone biosynthesis according to the KEGG database (https://www.genome.jp/kegg-bin/show_organism?org=T00066). Reactions are colored according to the essentiality of their corresponding genes.

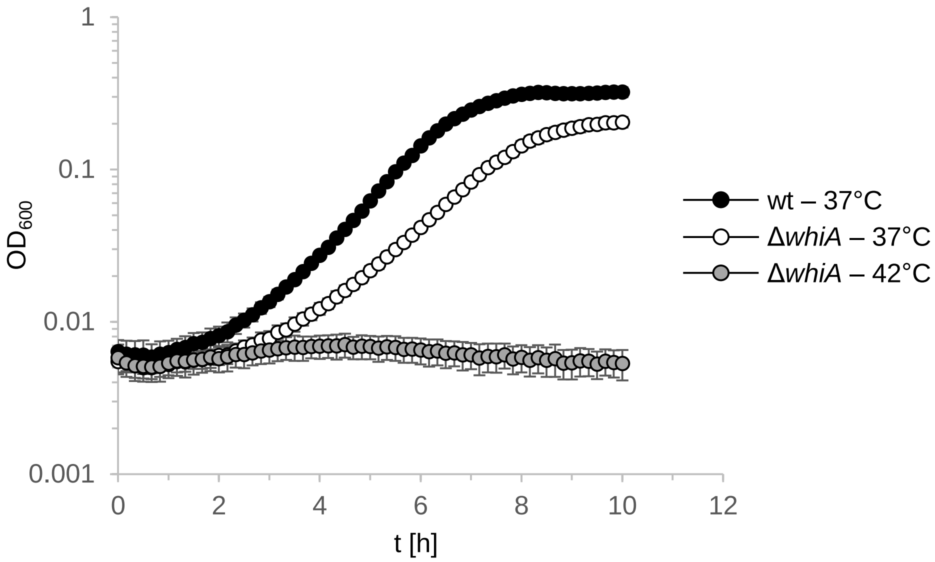


**Fig. S5:** Growth of a *L. monocytogenes* mutant lacking *whiA* (*lmo2472*)
Growth of *L. monocytogenes* strains EGD-e (wt) and LMS281(Δ*whiA*) in BHI broth at 37°C and 42°C. The experiment was performed with five technical replicates. Average values and standard deviations are shown.
