## Supplemental Table S2 for "*Listeria monocytogenes* gene essentiality under laboratory conditions and during macrophage infection"

**Tab. S2:** Genes with high PMS frequency in the *L. monocytogenes* population

| **locus_tag** | **gene** | **product** | **essentiality** | **PMS frequency (%)** |
|---|---|---|---|---|
| *lmo2760a* | | hypothetical protein | non-essential | 56.41 |
| *lmo0380* | | hypothetical protein | non-essential | 22.71 |
| *lmo0071* | | hypothetical protein | non-essential | 8.85 |
| *lmo1124* | | hypothetical protein | non-essential | 5.61 |
| *lmo0671* | | hypothetical protein, secreted | non-essential | 5.29 |
| *lmo0175* | | peptidoglycan-binding protein, LPXTG motif, | non-essential | 3.91 |
| *lmo0524* | | sulfate transporter | non-essential | 3.84 |
| *lmo1638* | | putative muramoyltetrapeptide carboxypeptidase | non-essential | 3.60 |
| *lmo0433* | *inlA* | internalin A | non-essential | 3.58 |
| *lmo2724* | | putative DNA binding 3-demethylubiquinone-9 3-methyltransferase domain protein | essential | 3.55 |
| *lmo2290* | | protein gp13 | non-essential | 2.64 |
| *lmo1412* | *flaR* | modulates DNA topology | non-essential | 2.56 |
| *lmo0117* | *lmaB* | antigen B | non-essential | 2.56 |
| *lmo1076* | *auto* | autolysin | non-essential | 2.53 |
| *lmo2809* | | hypothetical protein | non-essential | 2.43 |
| *lmo0148* | | hypothetical protein | non-essential | 2.34 |
| *lmo0586* | | hypothetical protein, CscA-like | non-essential | 2.14 |
| *lmo2733* | | PTS fructose transporter subunit IIABC | non-essential | 1.82 |
| *lmo1117* | | hypothetical protein | non-essential | 1.69 |
| *lmo0859* | | sugar ABC transporter substrate-binding protein | non-essential | 1.62 |
| *lmo0940* | | hypothetical protein | non-essential | 1.60 |
| *lmo2771* | | beta-glucosidase | non-essential | 1.35 |
| *lmo0395* | | blasticidin S-acetyltransferase | non-essential | 1.31 |
| *lmo1308* | | hypothetical protein | non-essential | 1.29 |
| *lmo0401* | | alpha-mannosidase | non-essential | 1.24 |
| *lmo2375* | | hypothetical protein | non-essential | 1.05 |
| *lmo2781* | | beta-glucosidase | non-essential | 1.05 |
| *lmo0660* | | transposase | non-essential | 1.00 |
